# Structural and mechanistic characterization of bifunctional heparan sulfate N-deacetylase-N-sulfotransferase 1

**DOI:** 10.1101/2023.08.30.555497

**Authors:** Courtney J. Mycroft-West, Sahar Abdelkarim, Helen M. E. Duyvesteyn, Neha S. Gandhi, Mark A. Skidmore, Raymond J. Owens, Liang Wu

## Abstract

Heparan sulfate (HS) polysaccharides are major constituents of the extracellular matrix, involved in myriad structural and signaling processes. Mature HS polysaccharides contain complex, non-templated patterns of sulfation and epimerization, which mediate interactions with diverse protein partners. Complex HS modifications form around initial clusters of glucosamine-N-sulfate (GlcNS) on nascent polysaccharide chains, but the mechanistic basis underpinning incorporation of the GlcNS modification itself into HS remains unclear. We have determined cryo-electron microscopy structures of human N-deacetylase-N-sulfotransferase (NDST)1, the bifunctional enzyme responsible for initial GlcNS modification of HS. Our structures reveal the architecture of both NDST1 deacetylase and sulfotransferase catalytic domains, alongside a previously unreported non-catalytic N-terminal domain. Surprisingly, the two catalytic domains of NDST1 adopt an unusual back-to-back topology that limits direct cooperativity. Binding analyses, aided by novel activity modulating nanobodies, suggest that sulfotransferase domain substrate anchoring initiates the NDST1 catalytic cycle, providing a plausible mechanism for cooperativity despite spatial domain separation. Our data shed light on key determinants of NDST1 activity, and describe tools to probe NDST1 function *in vitro* and *in vivo*.

## Introduction

Heparan sulfate (HS) is a ubiquitous and evolutionarily ancient class of polysaccharide produced throughout the metazoan lineage^1^. HS polysaccharides are abundant within the extracellular matrix (ECM) of multicellular organisms in the form of HS proteoglycans (HSPGs), wherein one or more polysaccharide chains are covalently linked to a membrane bound or pericellular core protein^2^. HSPGs in the ECM interact with myriad partners, and are essential regulators of physiological processes including growth factor signaling^3^, morphogen patterning^4^, adhesion^5^, endocytosis^6^, barrier filtration^7^ and host-pathogen binding^8–10^.

Compositionally, HS is a member of the glycosaminoglycan family - linear polysaccharides comprised of alternating hexosamine and uronic acid monosaccharides. The hexosamine units of HS are predominantly N-acetyl- or N-sulfo-glucosamine (GlcNAc, GlcNS respectively), and the uronic acids of HS are glucuronic or iduronic acid (GlcA, IdoA respectively). Each monosaccharide can be further modified by variable *O*-sulfation (**Figure S1a**)^11^, giving rise to diverse HS polysaccharide compositions that enable binding to multiple partners^12^. HS sequence complexity is crucial for its biological function, and is tightly regulated by cells in response to developmental cues, physiological status and external stimuli^13^.

HS biosynthesis begins with polymerization to produce unmodified [GlcNAc-GlcA]_n_ chains (hereafter heparosan) by the *EXT* family of glycosyltransferases^14–16^. Complexity is subsequently introduced by a series of modification reactions (**Figure S1b**), creating a modular polysaccharide structure in which ‘NS domains’, containing high proportions of IdoA and sulfated sugars, are interspersed by ‘NA domains’ more closely resembling heparosan. The biological functions of HS largely reside within its highly modified NS domains or NS/NA domain boundaries, which predominantly participate in protein interactions^2^.

Bifunctional N-deacetylase-N-sulfotransferase (NDST) enzymes are the first to act on nascent heparosan after its polymerization, converting GlcNAc sugars to GlcNS, *via* deacetylated glucosamine (GlcN) intermediates (**Figure S1c**). HS modification by NDSTs does not occur stochastically, but instead produces discrete GlcNS clusters of variable length^17,18^. Notably, these GlcNS clusters are themselves recognized by downstream enzymes, which work to generate further HS modifications, ultimately producing the highly sulfated and epimerized sequences characteristic of NS domains. The activity of NDST enzymes thus plays a central role in determining the overall locations and extents of modification within HS polysaccharide chains (**Figure 1a**)^13,19–21^.

**Figure 1.**
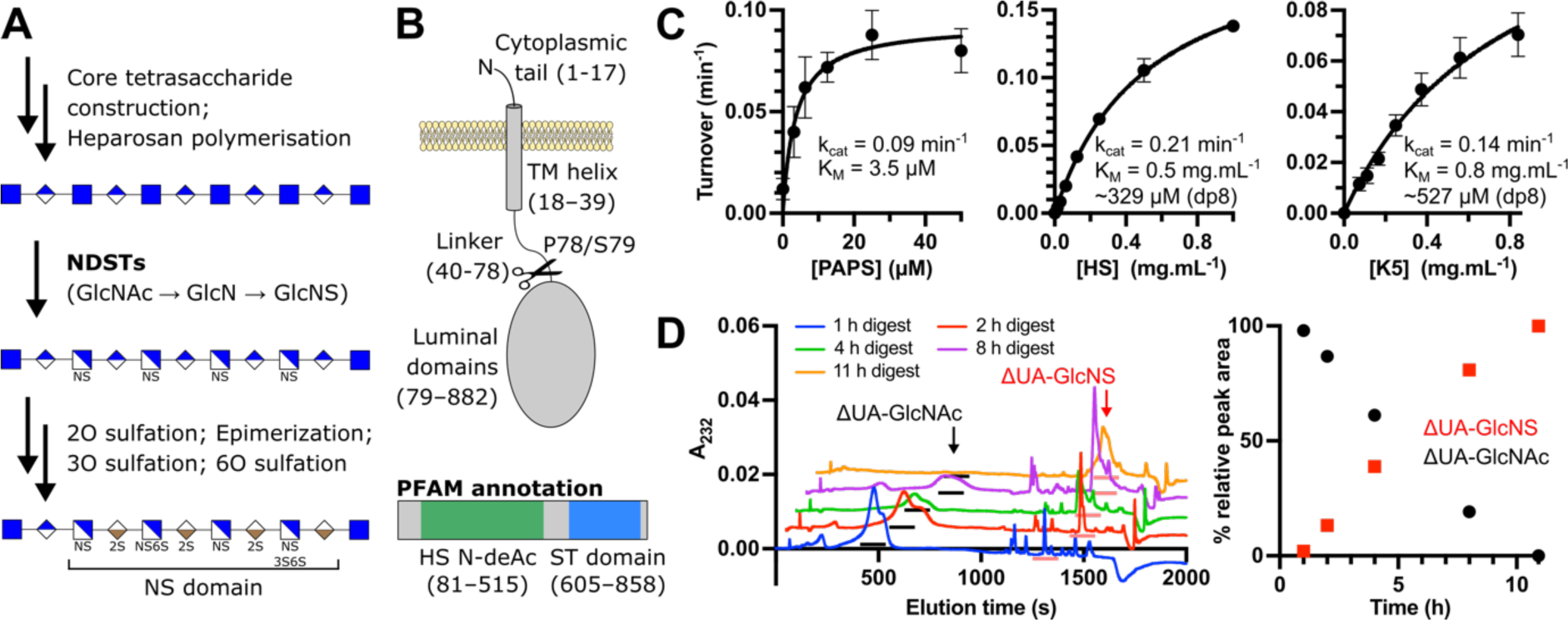
Activity profile of NDST1 (A) Sequential pathway for HS biosynthesis. The reaction catalyzed by NDST enzymes (conversion of GlcNAc to GlcNS via glucosamine) is highlighted in bold. GlcNS sugars produced by NDSTs are recognized and targeted by downstream modification enzymes to create highly-modified NS domains. (B) Schematic diagram of NDST1 on the Golgi membrane, and the site of truncation to make solubilized recombinant NDST1. PFAM annotation of NDST1 is shown, depicting predicted deacetylase (HS N-deAc) and sulfotransferase (ST) domains. (C) Pseudo-first order Michaelis-Menten kinetics for NDST1 with respect to sulfate donor substrate PAPS, and acceptor polysaccharide substrates HS and K5. (D) Disaccharide analysis of NDST1 activity against K5 polysaccharide, showing quantitative conversion of GlcNAc residues to GlcNS. Left – chromatographs of disaccharides generated from K5 after treatment with NDST1 for the indicated timepoints (staggered for clarity). Peaks corresponding to ΔUA-GlcNAc (black) and ΔUA-GlcNS (red) are annotated by arrows, and underscores below the peaks. Minor unannotated peaks are impurities from the reaction mixture. Right – Conversion of GlcNAc to GlcNS over time. Datapoints represent peak areas normalized to the total area of both ΔUA-GlcNAc and ΔUA-GlcNS peaks.

Humans express 4 members of the NDST family (NDST1–4). NDST1 and NDST2 are systemically distributed, with NDST1 dominant in most tissues, and NDST2 dominant in mast cells, which produce the HS analogue heparin^22,23^. NDST3 and NDST4 show more limited distribution, with both isoforms abundant in the brain^24–26^. Commensurate with their essential role in HS biosynthesis, dysregulation or mutations of NDSTs are associated with a spectrum of pathologies. Total loss of Ndst1 produces perinatal lethality in mice due to severe cerebral and craniofacial defects^27^, whilst partially inactivating *NDST1* mutations cause autosomal recessive intellectual disability in humans^28^. Changes to *NDST3* expression have been implicated in schizophrenia and bipolar disorder^29^, and loss of NDST4 has been suggested to be a prognostic marker for adverse colorectal cancers^30^.

Despite substantial interest, the molecular basis of NDST activity remains poorly understood. An X-ray crystal structure of the NDST1 sulfotransferase domain was reported in 1999 (PDB 1NST)^31^, providing some insight into the basis of sulfate transfer from 5’-phosphoadenosine-3’-phosphosulfate (PAPS) to deacetylated GlcN substrates. However, little is known about NDST deacetylase activity, and consequently, how the deacetylase and sulfotransferase activities of NDSTs are coordinated. Biochemical experiments have demonstrated reduced GlcNS formation and no clustering when HS oligosaccharides are modified by a mixture of truncated NDST1 deacetylase and sulfotransferase domains, strongly implying functional cooperativity in the full-length enzyme^18^. However, a lack of structural information has precluded analysis of the basis for such functional coupling to date.

Here, we have resolved high resolution cryogenic-electron microscopy (cryo-EM) structures of the major human NDST isoform NDST1. Structural characterization, aided by novel activity modulating nanobodies, reveal important determinants of NDST1 catalytic function, including active site architectures, and key residues and loops involved in substrate processing. In contrast to the known bifunctionality of NDST1, our structures clearly reveal a 3-domain enzyme architecture, with the sulfotransferase and deacetylase domains flanked by a previously unreported non-catalytic domain. Counterintuitively, we find that the deacetylase and sulfotransferase catalytic domains of NDST1 project in opposing directions, limiting possible mechanisms of coordination. Analysis of NDST1-HS binding suggests that initial anchoring interactions most likely occur at the sulfotransferase domain, despite deacetylation taking place first during the NDST1 catalytic cycle. Based on these observations, we propose a model for NDST1 bifunctionality that can operate even in the absence of direct domain-to-domain transfer.

## Results

### Production and characterization of recombinant NDST1

NDST1 is a Golgi lumen resident type-II transmembrane, which contains a single pass N-terminal helix that anchors the catalytic luminal portions of the enzyme to the Golgi membrane. To facilitate structural and biochemical characterization, we generated solubilized NDST1 truncated at P78/S79, removing the N-terminal helix and a linker region (**Figure 1b**). Solubilized recombinant NDST1 was expressed using a baculoviral expression system and purified to homogeneity *via* metal ion affinity, anion exchange, and size-exclusion chromatography.

We tested the *in vitro* enzymatic activity of solubilized NDST1 using a recently described coupled enzyme system, in which NDST1 activity is linked to the rat sulfotransferase Sult1A1, which catalytically regenerates sulfate donor PAPS from fluorogenic 4-methylumbelliferyl (4-MU) sulfate^32^ (**Figure S2a, b**). We carried out initial activity trials using Mg^2+^ supplemented in the assay buffer, to satisfy the requirement of NDST1 catalysis on divalent cations, which are necessary for deacetylation. Surprisingly, we also noted robust activity in the absence of divalent supplementation, which was inhibited by treatment with the chelators EDTA or dipicolinic acid (DPA) (**Figure S2c**). Inductively coupled plasma optical emission spectroscopy (ICP-OES) revealed that this was the result of residual NDST1 associated cations, primarily Ca^2+^ with trace Zn^2+^, carried over from protein purification. Further supplementation with Ca^2+^ did not improve activity, suggesting that carryover was stoichiometric (**Figure S2c, d**). Prior analysis by Dou *et al* suggests that NDST1 deacetylation is compatible with a range of metals, including Mg^2+^, Mn^2+^, Ca^2+^ and Co^2+^, with highest activity found in the presence of Ca^2+18^. We thus assigned the most likely identity of the metal cofactor in our NDST1 samples to be Ca^2+^.

Pseudo-first order activity constants were determined for NDST1 with respect to PAPS (k_cat_ = 0.09 min^-1^; K_M_ = 3.5 µM), as well as sulfate acceptors HS (k_cat_ = 0.21 min^-1^; K_M_ = 0.5 mg.mL^-1^) and *E. Coli* K5 – a capsular polysaccharide that shares the same repeating disaccharide motif as heparosan^33^ (k_cat_ = 0.14 min^-1^; K_M_ = 0.8 mg.mL^-1^; **Figure 1c**). NDST1 has previously been proposed to act upon 8-mer stretches of heparosan (MW 1519.3 Da)^18^, thus these K_M_ constants for HS and K5 approximate to ∼329 and ∼527 µM respectively.

To further confirm the activity of solubilized NDST1, we followed the conversion of GlcNAc to GlcNS in K5 polysaccharide by disaccharide analysis, employing exhaustive digestion by heparin lyase II from *P. heparinus*. NDST1 treatment of K5 resulted in time-dependent loss of Λ1UA-GlcNAc peaks, matched to increasing Λ1UA-GlcNS, clearly demonstrating enzymatic activity in a *bona fide* polysaccharide context (**Figure 1d**).

### Activity modulating nanobodies for NDST1

To aid mechanistic investigations of NDST1, we generated *de novo* nanobody (nAb) binders, which can provide insight by trapping conformational states relevant for function. A panel of NDST1 binding nAbs were isolated by primary immunization of a llama host with recombinant solubilized protein, followed by phage display bio-panning of the resulting immunized V_HH_ library over two rounds^34^.

Binding analysis using surface plasmon resonance (SPR) and biolayer interferometry (BLI) confirmed 5 NDST1 interacting nAbs. Binding affinities (K_D_s) spanned the nanomolar– micromolar range, in the rank order nAb7 > nAb13 **≈** nAb5 **≈** nAb6 >> nAb1, with rapid on/off kinetics in all cases (**Figure 2a**, **Figure S3**, **Table 1**). Additive SPR signals indicative of co-binding were detected for most pairwise combinations of nAbs, except nAb5+nAb7, nAb5+nAb13 and nAb7+nAb13, which displayed mutually exclusive binding (**Figure S4**).

**Figure 2.**
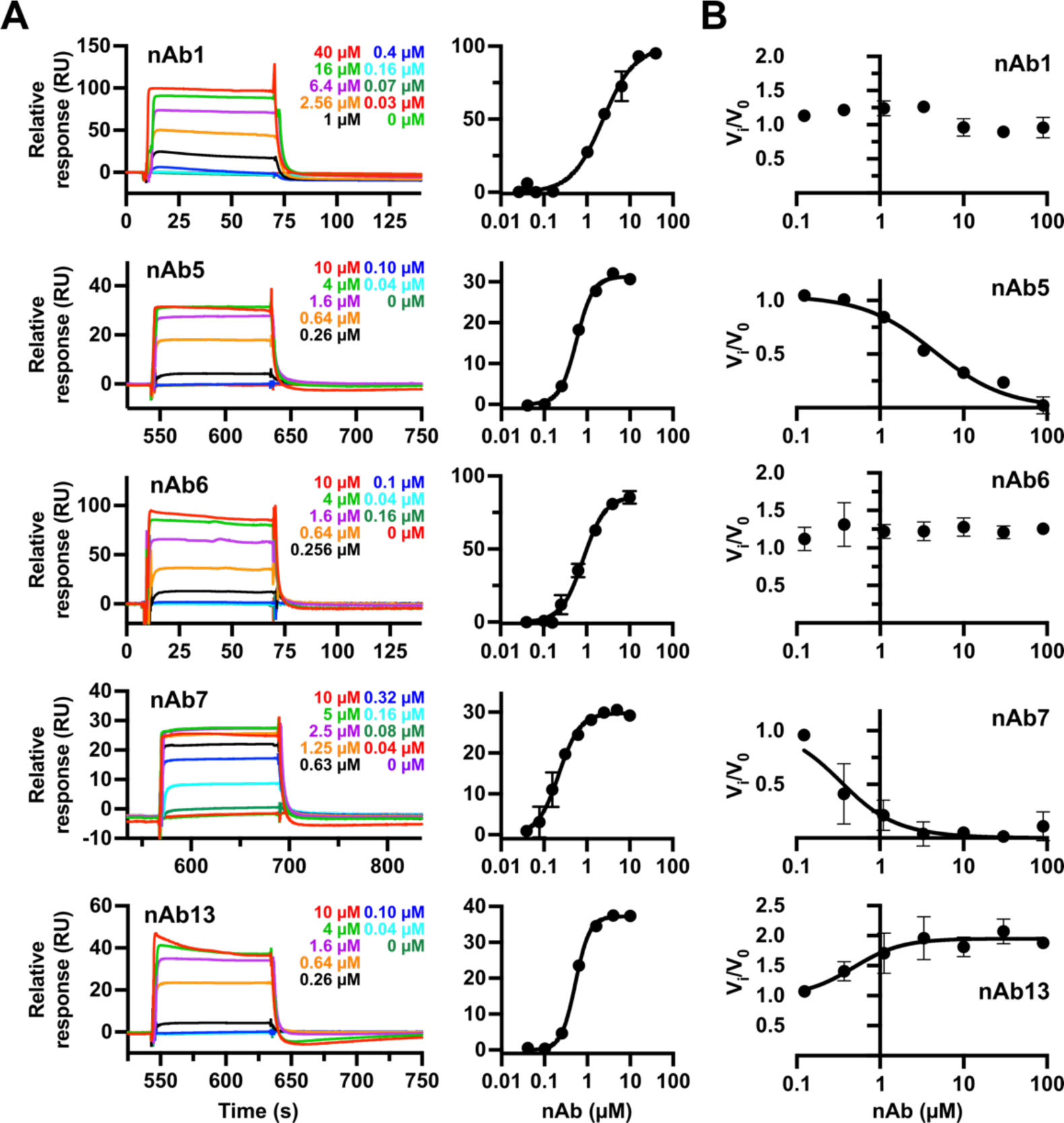
Biophysical characterization of anti-NDST1 nAbs (A) Representative SPR sensorgrams and steady state binding curves showing interaction of nAbs with surface bound NDST1. Datapoints on binding curves are mean ± s.d. (N = 3) (B) NDST1 activity modulation in the presence of nAbs. nAb5 and nAb7 are inhibitory, whilst nAb13 enhances NDST1 activity. Datapoints are mean ± s.d. (N = 3). Quantitated binding affinities and kinetic parameters are summarized in **Table 1**.

**Table 1.** NDST1 binding and activity modulation by nAbs, as measured by SPR, BLI and enzyme kinetics. N.D. not determinable due to experimental artefact.

| Nanobody | $K_D^*$ (SPR)/$\mu\text{M}$ | $K_D^*$ (BLI)/$\mu\text{M}$ | Activity modulation/$\mu\text{M}$ |
| --- | --- | --- | --- |
| nAb1 | 2.37 | 2.67 | > 100 |
| nAb5 | 0.57 | N.D. | 3.22 ($K_i$ inhibition) |
| nAb6 | 0.84 | 0.48 | > 100 |
| nAb7 | 0.22 | 0.12 | 0.15 ($K_i$ inhibition) |
| nAb13 | 0.53 | 0.24 | 0.48 ($\text{EC}_{50}$ enhancement) |

\*SPR and BLI affinities calculated from steady state binding curves

We also assessed nAbs for their ability to modulate NDST1 activity using the coupled enzyme assay. NDST1 inhibition was observed in the presence of nAb5 and nAb7, suggesting that these nAbs bind regions relevant for catalysis. Unexpectedly, we observed approximately 2-fold enhancement of NDST1 activity upon nAb13 binding, indicative of allosteric modulation that enhances catalytic turnover (**Figure 2b**, **Table 1**).

### Cryo-EM structure of bifunctional NDST1

To better understand the molecular basis for NDST1 activity, and how it can be modulated by nAb binding, we sought to resolve 3-dimensional structures of NDST1 and its complexes with nAb7 and nAb13 (**Figure S5**). A cryo-EM volume of NDST1 alone was calculated to a nominal resolution of 2.70 Å (Relion Au-FSC = 0.143), with the resulting map of sufficient quality to model of nearly all residues along the protein chain, using the AlphaFold2 model as a starting point for refinement (**Figure 3a**; **Figure S6**; **Table S1**).

**Figure 3.**
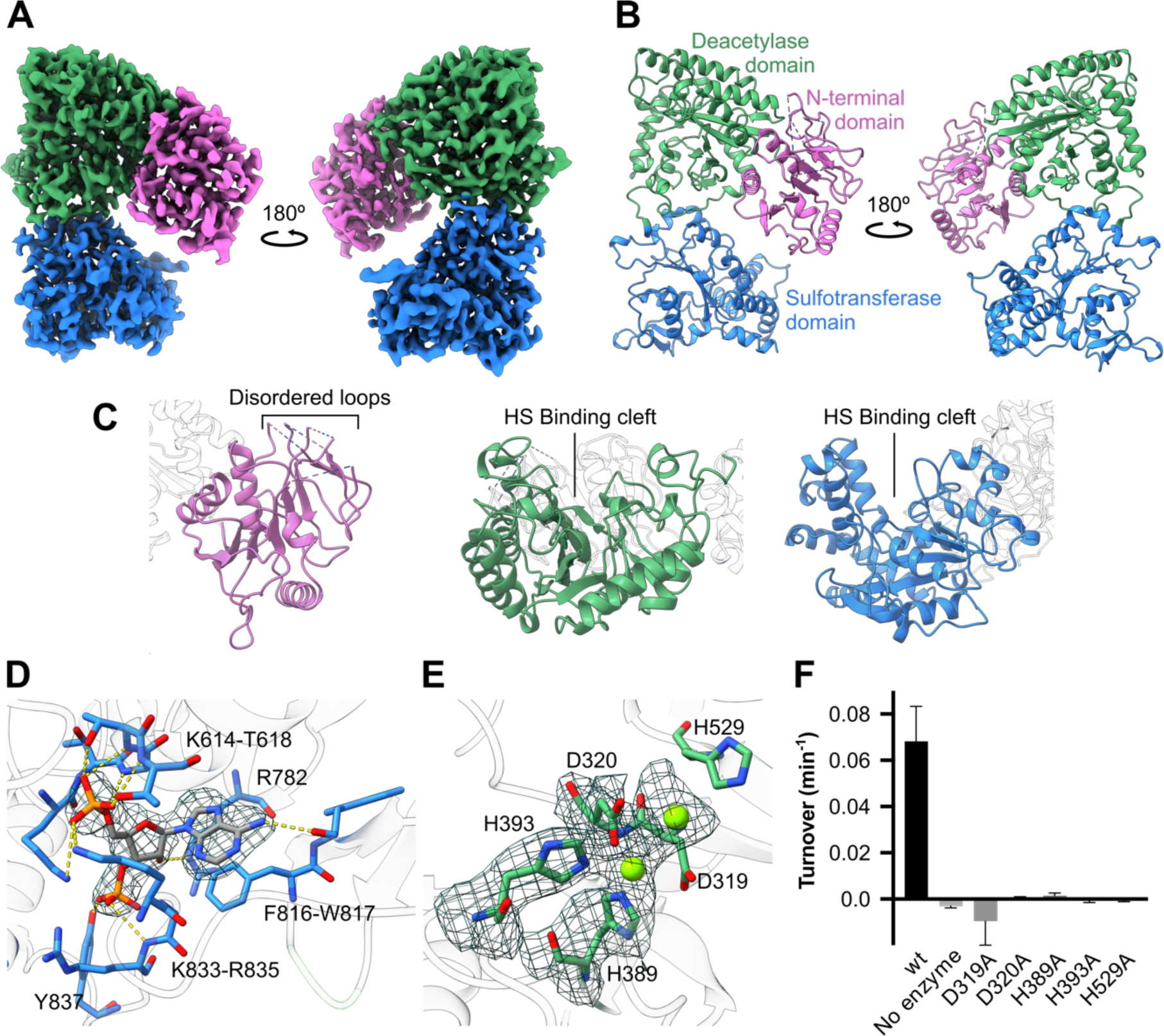
3-dimensional structure of NDST1. (A) Cryo-EM coulombic density map of NDST1 contoured to 12.30 σ, and colored by domain. (B) Ribbon model of NDST1 complex, colored by domain. (C) Close-up views of individual NDST1 domains, highlighting features of interest. Colors as in panel B. (D) Molecular architecture of the NDST1 sulfotransferase active site, showing interactions between bound PAP and neighboring amino acids. H-bonds are depicted by yellow dashes. Density contoured to 20.50 σ. (E) Molecular architecture of the NDST1 deacetylase active site, showing coordination of catalytic Ca^2+^ ions (green spheres) around the D320, H393, H389 triad. Density contoured to 15.40 σ. (F) Activity of NDST1 deacetylase domain mutants. Alanine mutation of residues at the catalytic center causes loss of activity.

NDST1 adopts an ‘elbow’ shaped monomeric structure comprised of 3 clearly resolved domains, in contrast to its widespread annotation as a bifunctional protein (**Figure 3b**; compare **Figure 1b**). The previously solved NDST1 sulfotransferase domain resides at the C-terminus of the protein (residues D602–R882), and our cryo-EM model agrees well with the crystal structure (PDB accession 1NST; RMSD 0.96 Å over 277 C⍺s; **Figure S7a**)^31^, displaying a well-defined binding cleft capable of accommodating HS polysaccharides (**Figure 3c**). We observed unambiguous density within the sulfotransferase active site corresponding to a molecule of 5’-phosphoadenosine-3’-phosphate (PAP; present in the cryo-EM sample), which binds NDST1 *via* H-bonds and salt bridges to K614–T618, R782, K833, R835, Y837, R782 and W817, as well as a π-stacking interaction from adenine to F816. The C828–D841 loop at the mouth of the PAP site partially occludes the ligand from solvent, suggesting a role as a stabilizing ‘lid’ that encloses PAP(S) after its engagement with the sulfotransferase domain (**Figure 3d**; **Figure S7b**). The positions of both PAP and the C828–D841 loop closely match the previously solved crystal structure.

Deacetylase activity was assigned to the central domain of NDST1 (L308–C601), based on close structural homology to the fungal galactosaminogalactan deacetylase Agd3 from *Aspergillus fumigatus* (PDB accession 6NWZ; 21.5% identity to NDST1 residues 1-601; **Figure S8a**, **b**)^35^. The deacetylase domain of NDST1 adopts a (β/α)_7_ barrel fold with a clear HS binding cleft (**Figure 3c**), and contains a His-His-Asp triad characteristic of metal dependent deacetylases^36^. Clear density, distinct from surrounding protein sidechains, was visible at the center of this triad, consistent with Ca^2+^ coordinated in a tridentate complex to H389, H393, D320. Further density within the deacetylase active site was tentatively modelled as a second Ca^2+^, consistent with the presence of multiple divalent ions in the active site of Agd3 and other deacetylases^35,37^ (**Figure 3e**; **Figure S8c**). To confirm the role of the deacetylase active site residues, we tested the activity of H389A, H393A and D320A mutants, as well as H529A and D319A – two nearby residues that likely function as general acid/base during deacetylation (see below). In all cases, mutation to alanine abrogated NDST1 turnover with minimal disruption to protein structure, demonstrating key roles for these sidechains in enzymatic processing (**Figure 3f**; **Figure S9**).

To our knowledge, the NDST1 N-terminal domain (NTD; residues 79–307) has not been previously reported. The NDST1 NTD adopts a Rossman-like fold, comprising a central layer of β-sheets flanked by α-helices on each side, although disorder prevented modelling of loops K171–S179, S191–D197, V215–W225 and K242–L262 (**Figure 3c**). Interestingly, a similar non-catalytic domain in Agd3 was suggested to function as a carbohydrate binding module, which extends the span of the substrate binding cleft^35^. Whilst the analogous cleft in NDST1 is occluded by the Q95–S106 helix (**Figure S8d**), inspection of the NDST1 surface does reveal a channel formed at the deacetylase and N-terminal domain interface, which extends the deacetylase cleft *via* a sharp turn (**Figure S8e**). This channel may guide long HS polysaccharides out of the deacetylase active site, and implicates a role for the NDST1 NTD in controlling enzyme-substrate interactions.

### Structures of NDST1-nAb complexes

NDST1-nAb7 and NDST1-nAb13 were isolated by preincubation followed by size-exclusion chromatography, and the resulting purified complexes analyzed by cryo-EM (**Figure S5**). Initial reconstructions of NDST1-nAb7 showed substantial disorder, consistent with unresolved heterogeneity, which was revealed by 3D variability analysis to correspond to a hinging motion around the NDST1 ‘elbow’ (**Supplementary Movie 1**). Further local refinements were thus carried out, focussed around either side of the ‘elbow joint’, yielding 2.41 Å partial maps (Au-FSC = 0.143) that were combined for model building purposes (**Figure S6**; **Table S1**). The resulting composite map was of sufficient quality to model both NDST1 and nAb7 proteins in detail, including the amino acid sidechains present at their interface.

Examination of the NDST1-nAb7 complex revealed nAb binding at the NDST1 deacetylase domain, across the ‘top’ of its substrate cleft, providing a clear steric rationale for inhibition (**Figure 4a, b**). nAb7 binding was predominantly mediated through its CDR3 loop, with H-bonds between D593_NDST1_–Y102_nAb7_, K465_NDST1_–Q113_nAb7_, R590_NDST1_–R99_nAb7_, K589_NDST1_–Y58_nAb7_ and salt bridges between D593_NDST1_–K101_nAb7_, E459_NDST1_–R99_nAb7_, K465_NDST1_–D99_nAb7_ and K465_NDST1_–D115_nAb7_ (**Figure 4c**; **Figure S10a**). The overall structure of NDST1 complexed to nAb7 was broadly similar to that of free NDST1, except for improved ordering of the D319– V332, C486–G512 and T528–L538 helical loops, likely due to interactions with the proximal nAb7 (**Figure S11**).

**Figure 4.**
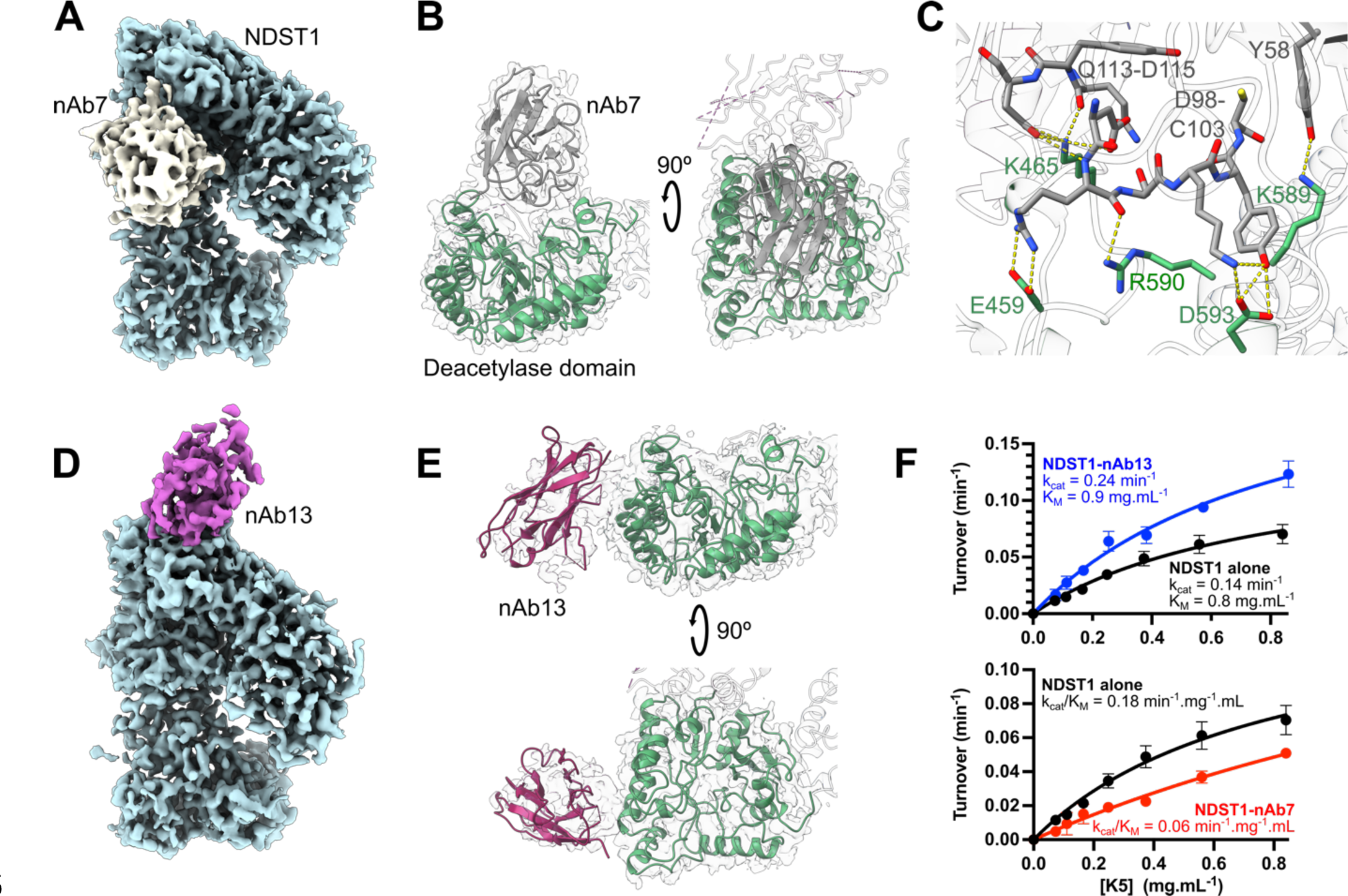
3-dimensional structures of NDST1 in complex with nAbs. (A) Coulombic density map of the NDST1-nAb7 complex contoured to 6.10 σ. Density for NDST1 is colored blue for clarity. Density for nAb7 is colored grey. (B) Ribbon depiction of the NDST1-nAb7 complex, showing occlusion of the NDST1 deacetylase cleft by the bound nAb. (C) Molecular interactions at the NDST1-nAb7 interface. H-bonds depicted by yellow dashes. Density for this interface is shown in **Figure S10a**. (D) Coulombic density map of the NDST1-nAb13 complex contoured to 6.90 σ. Density for nAb13 is colored purple. (E) Ribbon depiction of the NDST1-nAb13 complex, showing binding to the side of the NDST1 cleft. (F) Pseudo-first order Michaelis-Menten kinetics for NDST1, NDST1-nAb7 and NDST1-nAb13. nAb13 bound NDST1 shows enhanced turnover (higher k_cat_) compared to NDST1 alone. nAb7 bound NDST1 shows poorer substrate engagement (higher K_M_) compared to NDST1 alone.

A NDST1-nAb13 volume was also reconstructed following the same local refinement strategy as NDST1-nAb7, owing to a similar hinging movement (**Supplementary Movie 2**; **Figure S6**; **Table S1**). Consistent with its weaker binding, density for the nAb13 portion of the reconstructed map appeared disordered, and could only be modeled by rigid-body fitting of an all C⍺ nAb chain, derived from the nAb7 structure (**Figure 4d**).

nAb13 binds toward the ‘side’ of the deacetylase domain, in a region that does not obstruct the substrate cleft (**Figure 4e**). The rigid-body fitted nAb13 chain was found to lie within 6 Å of NDST1 residues K331, E333, K336 and R537, indicating these amino acids may be involved in binding (**Figure S10b**). As with nAb7, nAb13 binding improved ordering of the D319–V332, C486–G512 and T528–L538 loops, but did not otherwise alter NDST1 structure (**Figure S11**). No overlap was observed between the nAb7 and nAb13 binding sites, suggesting their inability to mutually bind NDST1 arises from allosteric effects (**Figure S4**; **Figure S12**).

We hypothesized that enhancement of NDST1 activity by nAb13 may be caused by ordering of the D319–V332, C486–G512 and T528–L538 loops, which together form one face of the deacetylase cleft, and likely contact HS during catalytic processing (**Figure S11**; **Figure S13**). Pre-organization of this interface by nAb13 would remove a substantial entropic penalty to HS engagement at the deacetylase domain, thereby enhancing the typically rate-limiting NDST1 deacetylation reaction^18^. Accordingly, Michaelis-Menten kinetics for NDST1-nAb13 with respect to K5 showed a ∼71% increased k_cat_ compared to NDST1 alone (0.24 min^-1^ vs 0.14 min^-1^), indicative of improved turnover efficiency. Conversely, NDST1-nAb7 displayed a ∼3x reduced k_cat_/K_M_ compared to NDST1 alone (0.06 min^-1^.mg^-1^.mL vs 0.18 min^-1^.mg^-1^.mL), primarily driven by increased K_M_, consistent with competitive inhibition of the deacetylase domain (**Figure 4f**).

### NDST1-HS interactions analyzed by molecular docking

Despite extensive trials, we were unable to resolve an experimental NDST1-HS complex by cryo-EM. We thus employed molecular docking to gain insights into NDST1 interactions with HS substrates, using GlycoTorch Vina^38^ to model the binding of unsulfated octasaccharide [GlcNAc-GlcA]_4_ (hereafter NS0) or di-N-sulfated octasaccharide [GlcNAc-GlcA]_2_-[GlcNS-GlcA]_2_ (NS2) to each catalytic domain of the nAb free NDST1 model.

Both NS0 and NS2 docked well into the NDST1 deacetylase cleft, placing their nonreducing termini towards the NTD, close to the channel formed at the deacetylase/NTD interface (**Figure 5a**; **Figure S8e**). The resulting contact surface incorporated residues from both the NDST1 deacetylase and N-terminal domains (**Figure S13a**). Notably, the second GlcNAc (from reducing end) of each octasaccharide was found to dock with its N-acetyl moiety close to a catalytic metal center, in a position ideally placed for deacetylation (**Figure 5b**). Whilst waters were not explicitly considered during docking, this docked ligand pose supports a mechanism (based on Agd3 homology) in which a metal coordinated water is deprotonated by the nearby sidechain of base D319, inducing nucleophilic attack upon the GlcNAc N-acetyl center. Breakdown of the resulting oxyanion intermediate would be aided by the nearby acid H529, leading to formation of GlcN with concomitant release of acetate. This mechanism is consistent with the inactivity of NDST1 deacetylase mutants (**Figure 3f**), as well as a previously proposed mechanisms for metal dependent deacetylases^35,36^.

**Figure 5.**
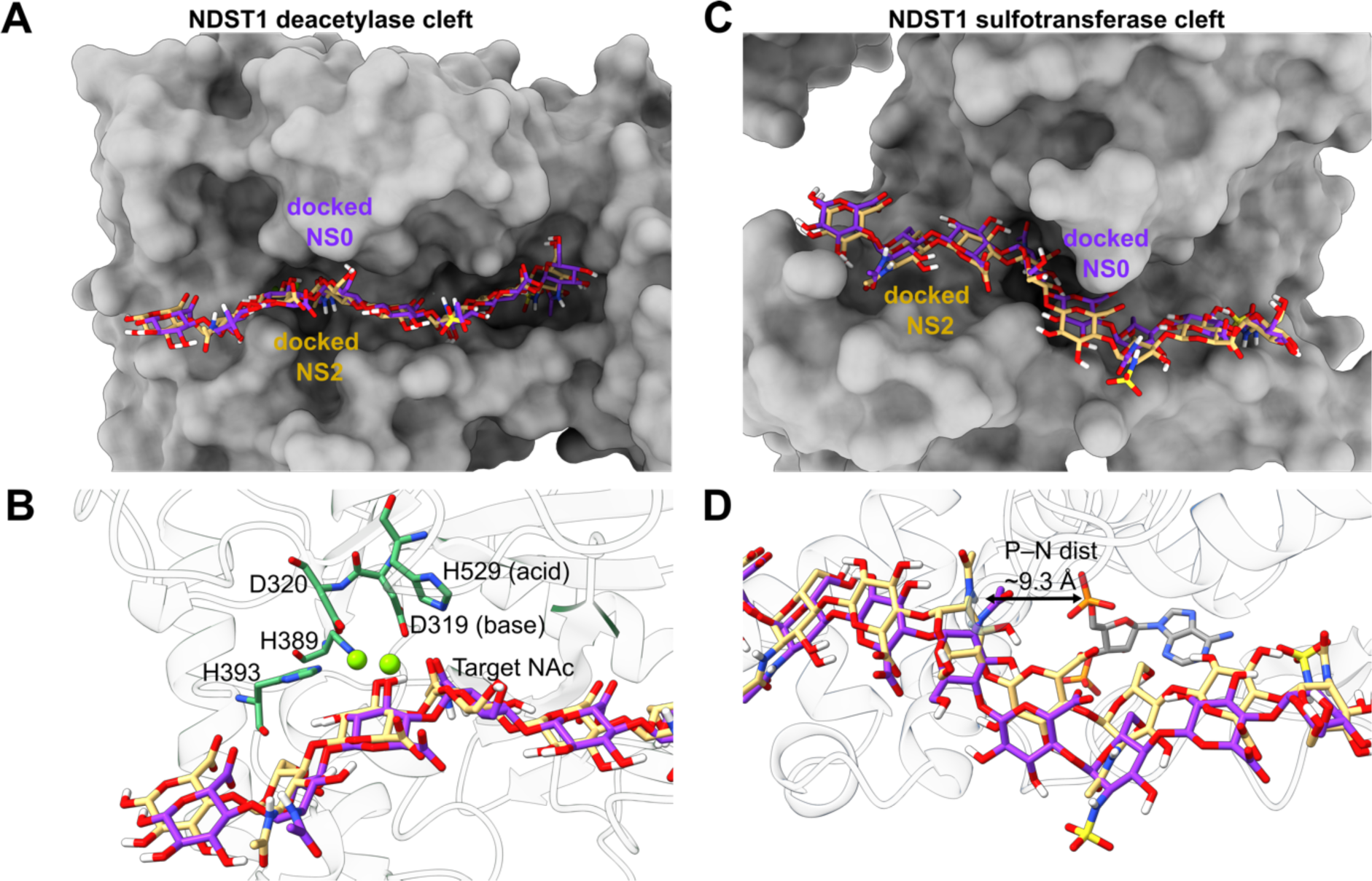
Computational docking of octasaccharides in active site clefts of NDST1. (A) Calculated docking poses for NS0 (purple) and NS2 (yellow) in the NDST1 deacetylase domain, showing similar carbohydrate trajectories within the binding cleft. (B) Close up of the deacetylase active site center, showing placement of the NAc substrate next to the catalytic amino acids and Ca^2+^ ions. (C) Calculated docking poses for NS0 (purple) and NS2 (yellow) in the NDST1 sulfotransferase domain. (D) Close up of the sulfotransferase active site center. The docked carbohydrates remain >9 Å distant from the bound PAP, possibly due to the lack of GlcN residues on both NS0 and NS2.

Docking in the NDST1 sulfotransferase domain showed both NS0 and NS2 closely tracking the active site cleft (**Figure 5c**; **Figure S13b**). The trajectory of both octasaccharides was similar to that observed for HS complexed with O-sulfotransferases HS3ST1 and HS3ST3 (PDB accessions 3UAN and 6XL8 respectively; **Figure S13c**), highlighting the conserved binding modes of these homologous enzymes. Although we did not observe close approach of docked NS0 and NS2 to the bound PAP ligand (**Figure 5d**; ∼9.3 Å between the PAP 5’ phosphorous and the closest GlcNAc nitrogen), it is plausible that deacetylated GlcN containing HS chains would be able to approach more closely to facilitate sulfate transfer from a sulfated PAPS donor.

### Sulfotransferase domain interactions initiate HS substrate engagement

Coordination between the deacetylase and sulfotransferase activities of NDST1 is essential for efficient HS modification and GlcNS cluster formation^14,16^. Unexpectedly, we found that the active sites of the NDST1 deacetylase and sulfotransferase domains project in opposing directions in the bifunctional enzyme, effectively precluding direct transfer of deacetylated HS intermediates from one domain to another (**Figure 6a**; **Figure S13d**).

**Figure 6.**
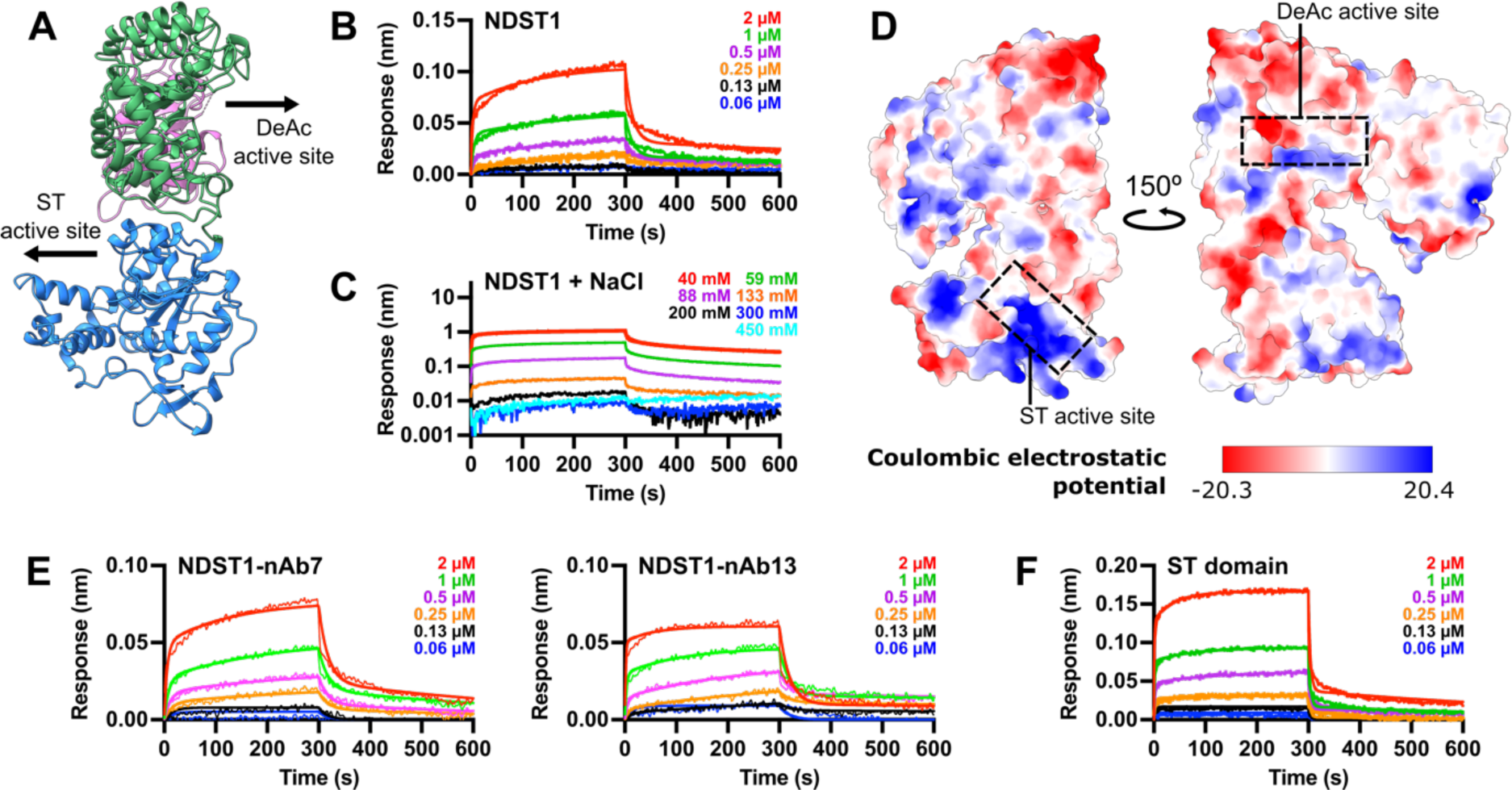
NDST1 domain organization and substrate binding analysis. (A) Ribbon diagram of the NDST1 structure, showing the opposing orientations of the deacetylase (DeAc) and sulfotransferase (ST) domain active sites. (B) BLI analysis of NDST1 binding to surface immobilized K5 polysaccharide, showing biphasic on/off kinetics. (C) NDST1 binding to K5 is abrogated at NaCl concentrations above ∼200 mM, suggesting a mainly electrostatic interaction. (D) Electrostatic surface potential of NDST1, calculated in ChimeraX. The sulfotransferase cleft contains a large positively charged region well-suited for binding HS. (E) NDST1-nAb7 and NDST1-nAb13 binding to surface immobilized K5. Similar nanomolar binding affinity is observed for the two NDST1-nAb complexes compared to NDST1 alone. (F) Truncated NDST sulfotransferase domain binding to surface immobilized K5. No substantial difference in affinity was observed compared to full length NDST1. All BLI binding constants are summarized in **Table 2**.

To determine possible mechanisms involved in domain cooperativity, we used BLI to measure the interaction of full length NDST1 to surface immobilized K5 polysaccharide. Binding of NDST1 to K5 could be well-modelled by a biphasic on/off model, with K_D_s of 234 nM and 679 nM respectively (**Figure 6b**; **Table 2**). Although biphasic interactions are not necessarily unusual for a multifunctional enzyme, we were intrigued by the substantial difference between the nanomolar K_D_s measured for direct NDST1-K5 binding, compared to the high-micromolar K_M_s determined for enzymatic NDST1-K5 processing (**Figure 1c**). Whilst K_D_ and K_M_ are not directly analogous, both constants quantitate interactions between proteins and ligands. A substantial difference between these values for NDST1-K5 suggested the presence of additional binding events that might take place alongside enzymatic processing.

**Table 2.**
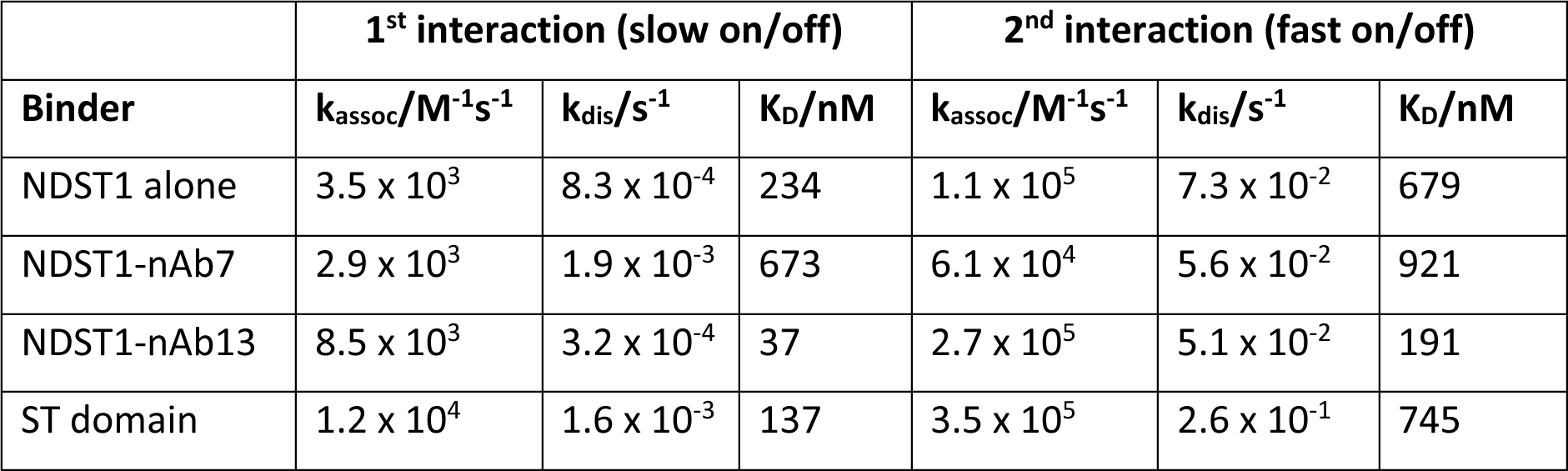
NDST1 binding affinities for K5. Full length protein, in the absence and presence of deacetylase domain binders nAb7 or nAb13, and truncated NDST1 sulfotransferase domain.

|  | 1<sup>st</sup> interaction (slow on/off) |  |  | 2<sup>nd</sup> interaction (fast on/off) |  |  |
| --- | --- | --- | --- | --- | --- | --- |
| Binder | $k_{\text{assoc}}/\text{M}^{-1}\text{s}^{-1}$ | $k_{\text{dis}}/\text{s}^{-1}$ | $K_{\text{D}}/\text{nM}$ | $k_{\text{assoc}}/\text{M}^{-1}\text{s}^{-1}$ | $k_{\text{dis}}/\text{s}^{-1}$ | $K_{\text{D}}/\text{nM}$ |
| NDST1 alone | $3.5 \times 10^3$ | $8.3 \times 10^{-4}$ | 234 | $1.1 \times 10^5$ | $7.3 \times 10^{-2}$ | 679 |
| NDST1-nAb7 | $2.9 \times 10^3$ | $1.9 \times 10^{-3}$ | 673 | $6.1 \times 10^4$ | $5.6 \times 10^{-2}$ | 921 |
| NDST1-nAb13 | $8.5 \times 10^3$ | $3.2 \times 10^{-4}$ | 37 | $2.7 \times 10^5$ | $5.1 \times 10^{-2}$ | 191 |
| ST domain | $1.2 \times 10^4$ | $1.6 \times 10^{-3}$ | 137 | $3.5 \times 10^5$ | $2.6 \times 10^{-1}$ | 745 |

HS-protein interfaces are typically characterized by charge-based interactions between anionic polysaccharides and cationic surfaces on proteins. Nascent heparosan polysaccharides lack sulfates, but still possess considerable negative charge due to their repetitive carboxylate moieties. We assessed the binding of NDST1 to K5 polysaccharide in the presence of increasing concentrations of NaCl, to disrupt charge-based interactions. NaCl concentrations above ∼200 mM essentially abrogated all binding, consistent with an electrostatic interface between NDST1 and HS, and in line with functional reports that NaCl strongly inhibits NDST1 processing^39^ (**Figure 6c**).

To identify regions on NDST1 that may bind HS *via* charge, we computed an electrostatic surface for NDST1 using the nAb free structure, which revealed substantial cationic potential around the sulfotransferase cleft, contrasting with neutral to net-anionic potential at the deacetylase cleft (**Figure 6d**). Whilst such charge-based considerations suggest initial HS binding at the sulfotransferase domain, deacetylation is known to occur first during the NDST1 catalytic cycle (**Figure S1c**). We thus reassessed NDST1-K5 binding in the presence of 10 µM nAb7 or nAb13 (>> 10x K_D_ for both nAbs), which bind to different epitopes of the NDST1 deacetylase domain, with nAb7 also binding in a fashion that occludes polysaccharide. As with full length NDST1, nanomolar K5 affinities were measured for both NDST1-nAb7 and NDST1-nAb13, thereby effectively ruling out the NDST1 deacetylase cleft as a site of initial substrate binding (**Figure 6e**; **Table 2**). Finally, we tested the binding of truncated NDST1 sulfotransferase domain alone to K5 polysaccharide. The NDST1 sulfotransferase domain displayed essentially equivalent nanomolar affinity for K5 as compared to full length enzyme, further reinforcing the primacy of this domain in driving NDST1 polysaccharide interactions (**Figure 6f**; **Table 2**).

Taken together, our data indicate that initial, high-affinity NDST1-HS binding likely takes place at the sulfotransferase domain, despite deacetylation being the first step of the enzymatic processing cycle.

## Discussion

Mammalian HS biosynthesis is a complex multistep process, which involves the coordinated activity of multiple enzymes along the ER-Golgi pathway. NDSTs are key players in the early stages of HS maturation, which convert GlcNAc sugars in nascent heparosan to GlcNS. Clusters of GlcNS laid down by NDSTs play a major role in directing downstream HS processing, thus understanding NDST function fundamentally informs upon how cells create the complex HS sequence modifications that are required by biology.

Here, we carried out detailed structural and functional characterization of NDST1, the dominant human NDST isoform, revealing a 3-domain enzyme, which possesses a previously unreported NTD in additional to its catalytic deacetylase and sulfotransferase domains. The N-terminal and deacetylase domains of NDST1 have not previously been experimentally resolved in detail. Computational docking of NDST1 with HS octasaccharides suggests that the NTD may play a role in polysaccharide engagement, *via* the formation of a substrate cleft adjacent to the deacetylase domain. The deacetylase domain of NDST1 is characterized by a His-His-Asp triad, typical of metal dependent sugar deacetylases. Herein, we have assigned the presence of catalytic Ca^2+^ coordinated by this triad, based on its high affinity carry through from protein purification. However, NDST1 has also been shown to display activity in the presence of other divalent ions, including Mn^2+^, Mg^2+^ and Co^2+ 18^. It is possible that physiological NDST1 within the Golgi apparatus, which is enriched in many divalent cations^40^, may employ a variety of metal ions to catalyze deacetylation.

Unusually, the active sites of the NDST1 deacetylase and sulfotransferase catalytic domains project in opposite directions, precluding a simple substrate transfer mechanism for bifunctionality. Our data suggest that initial NDST1-HS binding occurs at the sulfotransferase domain *via* an electrostatic interface, despite deacetylation being the first step of the NDST1 catalytic cycle. This initial sulfotransferase binding occurs with higher affinity than that required for enzymatic conversion, and may help NDST1 accumulate on nascent polysaccharides prior to catalytic processing. Initial sulfotransferase binding also provides a ready mechanism for bifunctionality, which obviates the need for direct domain to domain substrate transfer. Putatively, deacetylation of HS polysaccharides anchored to the NDST1 sulfotransferase domain would produce GlcN intermediates that remain bound to the enzyme. Such intermediates, already located at the NDST1 sulfotransferase domain, would consequently be ideally positioned for immediate sulfation to produce GlcNS. Because initial NDST1-HS interactions are predominantly electrostatic, they may also bind newly formed GlcNS preferentially, further raising the intriguing possibility of a molecular ‘ratcheting’ mechanism that can lead to the formation of GlcNS clusters (**Figure S14**). Consistent with this, we note that NDST1 processing of HS, which contains some N-sulfation, shows improved K_M_ and k_cat_ compared to its processing of K5, which entirely lacks N-sulfation (**Figure 1c**). Given the constraints on HS binding imposed by the NDST1 domain architecture (**Figure 6a**), it appears likely that deacetylation would require a second enzyme molecule. Although our data show no evidence for the presence of stable NDST1 dimers, implying that deacetylation may instead occur *via* transient dimerization events, recent observations support the fact that NDST1 dimerization is fully compatible with activity^41^. We also do not discount the possibility of more stable multimeric states within intracellular contexts, formed by the organization of enzymes on Golgi membranes. Detailed analysis of NDST1-HS interactions across multiple polysaccharide sulfation states will be required to verify the consequences of different NDST1 substrate binding modes and oligomeric states.

One final aspect of NDST1 function we have not directly addressed is the formation of free GlcN residues, which typically comprise ∼0.7–4% of HS sequences^42^. These rare GlcN residues may arise from independent NDST1 deacetylase domain activity upon a HS chain, or alternatively, a catalytic cycle stalled at the sulfotransferase step (*e.g.* due to lack of PAPS), resulting in premature release of deacetylated intermediate. Both possibilities are likely to be rare, given the marginal K_M_ of NDST1 deacetylase function (**Fig 1c**), and the abundance of sulfation reactions within cells requiring a stable pool of PAPS, respectively. Such mechanistic rarity is consistent with the paucity of free GlcN in mature HS.

Important to our characterization of NDST1 was the generation of a novel panel of nAb binders, which aided our analysis of NDST1-HS interactions, and shed light on allosteric factors that influence NDST1 catalytic activity. Beyond their clear utility for functional analysis, our nAbs provide a potential toolkit for functional studies of NDST1 biology *e.g.* as probes to track enzymes *in situ*^43^. Notably, we have isolated nAbs that both inhibit (nAb7), or enhance (nAb13) NDST1 activity. Given the paucity of high-quality tools to perturbate HS biosynthesis in cells, these nAbs may represent useful, genetically encodable modules to up- or downregulate HS modifications in a reversible manner. *In situ* use may require higher binding affinities than the mid-nM potencies currently possessed by nAb7 and nAb13. Thus affinity maturation steps may form part of future efforts to develop HS modulating nAb binders^44^.

With our characterization of bifunctional NDST1, all enzyme families involved in the mammalian HS biosynthesis pathway (linker biosynthesis, polymerization, N-sulfation, epimerization, 2O-, 3O- and 6O-sulfation) now have at least one experimentally resolved representative^14,45–51^. The ‘GAGosome’ hypothesis, first proposed over 20 years ago, posits the existence of a large macromolecular assembly of biosynthesis enzymes tasked with constructing HS in a coordinated fashion^17^. Although some HS biosynthesis enzyme interactions have been inferred by coimmunoprecipitation or immunofluorescence, the only multimers to have been directly visualized remain those between the obligate dimeric EXT enzymes^14–16^. Our observation of initial high-affinity binding interactions between NDST1 and HS hints at an alternative model for the ‘GAGosome’, in which HS itself may induce formation of complexes that transiently assemble around newly synthesized polysaccharide chains. Detailed analysis of individual HS biosynthesis enzymes such as NDST1 provides a vital basis for understanding mechanisms of cooperativity within complex multicomponent systems such as the ‘GAGosome’, and will underpin future studies of HS biosynthesis within *in situ* cellular contexts.

## Methods

Detailed methods, **Figures S1-S14**,**Table S1** and **Supplemental Videos 1-2** can be found online in the Supplemental Information.

## Data availability

Cryo-EM coordinates have been deposited in the PDB and EMDB under accession codes 8CCY and EMD-16564 (nAb free NDST1), 8CD0, EMD-16627, EMD-16629, EMD-16565 (NDST1-nAb7), 8CHS, EMD-16662, EMD-16663, EMD-16664 (NDST1-nAb13).

## Contributions

LW and CJMW conceived and designed experiments. LW and CJMW carried out all protein expression and purification. CJMW carried out polysaccharide production and purification, with assistance from MAS. CJMW and SA generated nanobodies, under supervision from RJO. CJMW carried out all biochemical and biophysical analyses. CJMW, HMED and LW carried out cryo-EM sample preparation, data collection and data analysis. NSG carried out molecular docking simulations. LW wrote the manuscript with input from all authors.

## Acknowledgements

This work was supported by the Rosalind Franklin Institute, with funding delivery partner the Engineering and Physical Sciences Research Council (EPSRC) UK, and a Biotechnology and Biological Sciences Research Council UK (BBSRC) grant for Nanobody Discovery (BB/V018523/1). We thank Jiandong Huo for assistance with phage display library construction, and Mark Arrowsmith at the University of Keele School of Pharmacy and Bioengineering for assistance with ICP-OES. Electron microscopy provision was provided *via* the Rosalind Franklin institute, and through the Oxford Particle Imaging Centre (OPIC) facility, a UK Instruct-ERIC Centre founded by a Wellcome JIF award (060208/Z/00/Z) and supported by a Wellcome equipment grant (093305/Z/10/Z). Computation for cryo-EM analysis was performed using the Oxford Biomedical Research Computing (BMRC) facility, a joint development between the Wellcome Centre for Human Genetics and the Big Data Institute supported by Health Data Research UK and the NIHR Oxford Biomedical Research Centre. Financial support for HMED was provided by the Wellcome Trust Core Award Grant Number 203141/Z/16/Z. MS acknowledges support from the Biotechnology and Biological Sciences Research Council, UK (BB/L023717/1 & BB/X011739/1) and the Engineering and Physical Sciences Research Council, UK (EP/X019179/1). LW is supported by a Sir Henry Dale Fellowship, jointly funded by the Wellcome Trust and the Royal Society (218579/Z/19/Z). NSG is supported by QUT Accelerate Fellowship. Computational (and/or data visualization) resources and services used in this work were provided by the eResearch Office, Queensland University of Technology, Brisbane, Australia. NSG would also like to acknowledge the National Computational Infrastructure (NCI), Australia for providing the high-performance computing (HPC) facility to carry out molecular docking work.

## References

1. Esko, J.D. & Lindahl, U. Molecular diversity of heparan sulfate. J Clin Invest 108, 169–73 (2001).

2. Sarrazin, S., Lamanna, W.C. & Esko, J.D. Heparan sulfate proteoglycans. Cold Spring Harb Perspect Biol 3(2011).

3. Pellegrini, L. Role of heparan sulfate in fibroblast growth factor signalling: a structural view. Curr Opin Struct Biol 11, 629–34 (2001).

4. Yan, D. & Lin, X. Shaping morphogen gradients by proteoglycans. Cold Spring Harb Perspect Biol 1, a002493 (2009).

5. Lim, H.C., Multhaupt, H.A. & Couchman, J.R. Cell surface heparan sulfate proteoglycans control adhesion and invasion of breast carcinoma cells. Mol Cancer 14, 15 (2015).

6. Christianson, H.C. & Belting, M. Heparan sulfate proteoglycan as a cell-surface endocytosis receptor. Matrix Biol 35, 51–5 (2014).

7. Bode, L. et al. Heparan sulfate and syndecan-1 are essential in maintaining murine and human intestinal epithelial barrier function. J Clin Invest 118, 229–38 (2008).

8. Clausen, T.M. et al. SARS-CoV-2 Infection Depends on Cellular Heparan Sulfate and ACE2. Cell 183, 1043–1057 e15 (2020).

9. Mycroft-West, C.J. et al. Heparin Inhibits Cellular Invasion by SARS-CoV-2: Structural Dependence of the Interaction of the Spike S1 Receptor-Binding Domain with Heparin. Thromb Haemost 120, 1700–1715 (2020).

10. Shukla, D. & Spear, P.G. Herpesviruses and heparan sulfate: an intimate relationship in aid of viral entry. J Clin Invest 108, 503–10 (2001).

11. Rabenstein, D.L. Heparin and heparan sulfate: structure and function. Nat Prod Rep 19, 312–31 (2002).

12. Meneghetti, M.C. et al. Heparan sulfate and heparin interactions with proteins. J R Soc Interface 12, 0589 (2015).

13. Kreuger, J. & Kjellen, L. Heparan sulfate biosynthesis: regulation and variability. J Histochem Cytochem 60, 898–907 (2012).

14. Leisico, F. et al. Structure of the human heparan sulfate polymerase complex EXT1-EXT2. Nature Communications 13, 7110 (2022).

15. Wilson, L.F.L. et al. The structure of EXTL3 helps to explain the different roles of bi-domain exostosins in heparan sulfate synthesis. Nature Communications 13(2022).

16. Li, H. et al. Structural basis for heparan sulfate co-polymerase action by the EXT1-2 complex. Nat Chem Biol (2023).

17. Esko, J.D. & Selleck, S.B. Order out of chaos: assembly of ligand binding sites in heparan sulfate. Annu Rev Biochem 71, 435–71 (2002).

18. Dou, W., Xu, Y., Pagadala, V., Pedersen, L.C. & Liu, J. Role of Deacetylase Activity of N-Deacetylase/N-Sulfotransferase 1 in Forming N-Sulfated Domain in Heparan Sulfate. J Biol Chem 290, 20427–37 (2015).

19. Annaval, T. et al. Heparan Sulfate Proteoglycans Biosynthesis and Post Synthesis Mechanisms Combine Few Enzymes and Few Core Proteins to Generate Extensive Structural and Functional Diversity. Molecules 25(2020).

20. Sheng, J., Liu, R., Xu, Y. & Liu, J. The dominating role of N-deacetylase/N-sulfotransferase 1 in forming domain structures in heparan sulfate. J Biol Chem 286, 19768–76 (2011).

21. Carlsson, P., Presto, J., Spillmann, D., Lindahl, U. & Kjellen, L. Heparin/heparan sulfate biosynthesis: processive formation of N-sulfated domains. J Biol Chem 283, 20008–14 (2008).

22. Kjellen, L. Glucosaminyl N-deacetylase/N-sulphotransferases in heparan sulphate biosynthesis and biology. Biochem Soc Trans 31, 340–2 (2003).

23. Pikas, D.S., Eriksson, I. & Kjellen, L. Overexpression of different isoforms of glucosaminyl N-deacetylase/N-sulfotransferase results in distinct heparan sulfate N-sulfation patterns. Biochemistry 39, 4552–8 (2000).

24. Aikawa, J. & Esko, J.D. Molecular cloning and expression of a third member of the heparan sulfate/heparin GlcNAc N-deacetylase/ N-sulfotransferase family. J Biol Chem 274, 2690–5 (1999).

25. Aikawa, J., Grobe, K., Tsujimoto, M. & Esko, J.D. Multiple isozymes of heparan sulfate/heparin GlcNAc N-deacetylase/GlcN N-sulfotransferase. Structure and activity of the fourth member, NDST4. J Biol Chem 276, 5876–82 (2001).

26. Pallerla, S.R. et al. Altered heparan sulfate structure in mice with deleted NDST3 gene function. J Biol Chem 283, 16885–94 (2008).

27. Grobe, K. et al. Cerebral hypoplasia and craniofacial defects in mice lacking heparan sulfate Ndst1 gene function. Development 132, 3777–86 (2005).

28. Reuter, M.S. et al. NDST1 missense mutations in autosomal recessive intellectual disability. Am J Med Genet A 164A, 2753–63 (2014).

29. Lencz, T. et al. Genome-wide association study implicates NDST3 in schizophrenia and bipolar disorder. Nat Commun 4, 2739 (2013).

30. Tzeng, S.T. et al. NDST4 is a novel candidate tumor suppressor gene at chromosome 4q26 and its genetic loss predicts adverse prognosis in colorectal cancer. PLoS One 8, e67040 (2013).

31. Kakuta, Y., Sueyoshi, T., Negishi, M. & Pedersen, L.C. Crystal structure of the sulfotransferase domain of human heparan sulfate N-deacetylase/ N-sulfotransferase 1. J Biol Chem 274, 10673–6 (1999).

32. Atienza, J., Tkachyova, I., Tropak, M., Fan, X. & Schulze, A. Fluorometric coupled enzyme assay for N-sulfotransferase activity of N-deacetylase/N-sulfotransferase (NDST). Glycobiology 31, 1093–1101 (2021).

33. Ly, M. et al. Analysis of E. coli K5 capsular polysaccharide heparosan. Anal Bioanal Chem 399, 737–45 (2011).

34. Pardon, E. et al. A general protocol for the generation of Nanobodies for structural biology. Nat Protoc 9, 674–93 (2014).

35. Bamford, N.C. et al. Structural and biochemical characterization of the exopolysaccharide deacetylase Agd3 required for Aspergillus fumigatus biofilm formation. Nat Commun 11, 2450 (2020).

36. Burger, M. & Chory, J. Structural and chemical biology of deacetylases for carbohydrates, proteins, small molecules and histones. Communications Biology 1(2018).

37. Ahangar, M.S. et al. Structural and functional determination of homologs of the Mycobacterium tuberculosis N-acetylglucosamine-6-phosphate deacetylase (NagA). J Biol Chem 293, 9770–9783 (2018).

38. Boittier, E.D., Burns, J.M., Gandhi, N.S. & Ferro, V. GlycoTorch Vina: Docking Designed and Tested for Glycosaminoglycans. J Chem Inf Model 60, 6328–6343 (2020).

39. Wei, Z., Swiedler, S.J., Ishihara, M., Orellana, A. & Hirschberg, C.B. A single protein catalyzes both N-deacetylation and N-sulfation during the biosynthesis of heparan sulfate. Proc Natl Acad Sci U S A 90, 3885–8 (1993).

40. Kellokumpu, S. Golgi pH, Ion and Redox Homeostasis: How Much Do They Really Matter? Front Cell Dev Biol 7, 93 (2019).

41. Vallet, S.D., et al. Functional and structural insights into human N-deacetylase/N-sulfotransferase activities. Proteoglycan Research 1, e8 (2023).

42. Westling, C. & Lindahl, U. Location of N-unsubstituted glucosamine residues in heparan sulfate. J Biol Chem 277, 49247–55 (2002).

43. Traenkle, B. & Rothbauer, U. Under the Microscope: Single-Domain Antibodies for Live-Cell Imaging and Super-Resolution Microscopy. Front Immunol 8, 1030 (2017).

44. Doria-Rose, N.A. & Joyce, M.G. Strategies to guide the antibody affinity maturation process. Curr Opin Virol 11, 137–47 (2015).

45. Xu, Y. et al. Structure Based Substrate Specificity Analysis of Heparan Sulfate 6-O-Sulfotransferases. ACS Chem Biol 12, 73–82 (2017).

46. Wander, R. et al. Structural and substrate specificity analysis of 3-O-sulfotransferase isoform 5 to synthesize heparan sulfate. ACS Catal 11, 14956–14966 (2021).

47. Tsutsui, Y., Ramakrishnan, B. & Qasba, P.K. Crystal structures of beta-1,4-galactosyltransferase 7 enzyme reveal conformational changes and substrate binding. J Biol Chem 288, 31963–70 (2013).

48. Pedersen, L.C., et al. Heparan/chondroitin sulfate biosynthesis. Structure and mechanism of human glucuronyltransferase I. J Biol Chem 275, 34580–5 (2000).

49. Liu, C. et al. Molecular mechanism of substrate specificity for heparan sulfate 2-O-sulfotransferase. J Biol Chem 289, 13407–18 (2014).

50. Debarnot, C. et al. Substrate binding mode and catalytic mechanism of human heparan sulfate d-glucuronyl C5 epimerase. Proc Natl Acad Sci U S A 116, 6760–6765 (2019).

51. Briggs, D.C. & Hohenester, E. Structural Basis for the Initiation of Glycosaminoglycan Biosynthesis by Human Xylosyltransferase 1. Structure 26, 801–809 e3 (2018).

