## Supplemental Information for "Structural and mechanistic characterization of bifunctional heparan sulfate N-deacetylase-N-sulfotransferase 1"

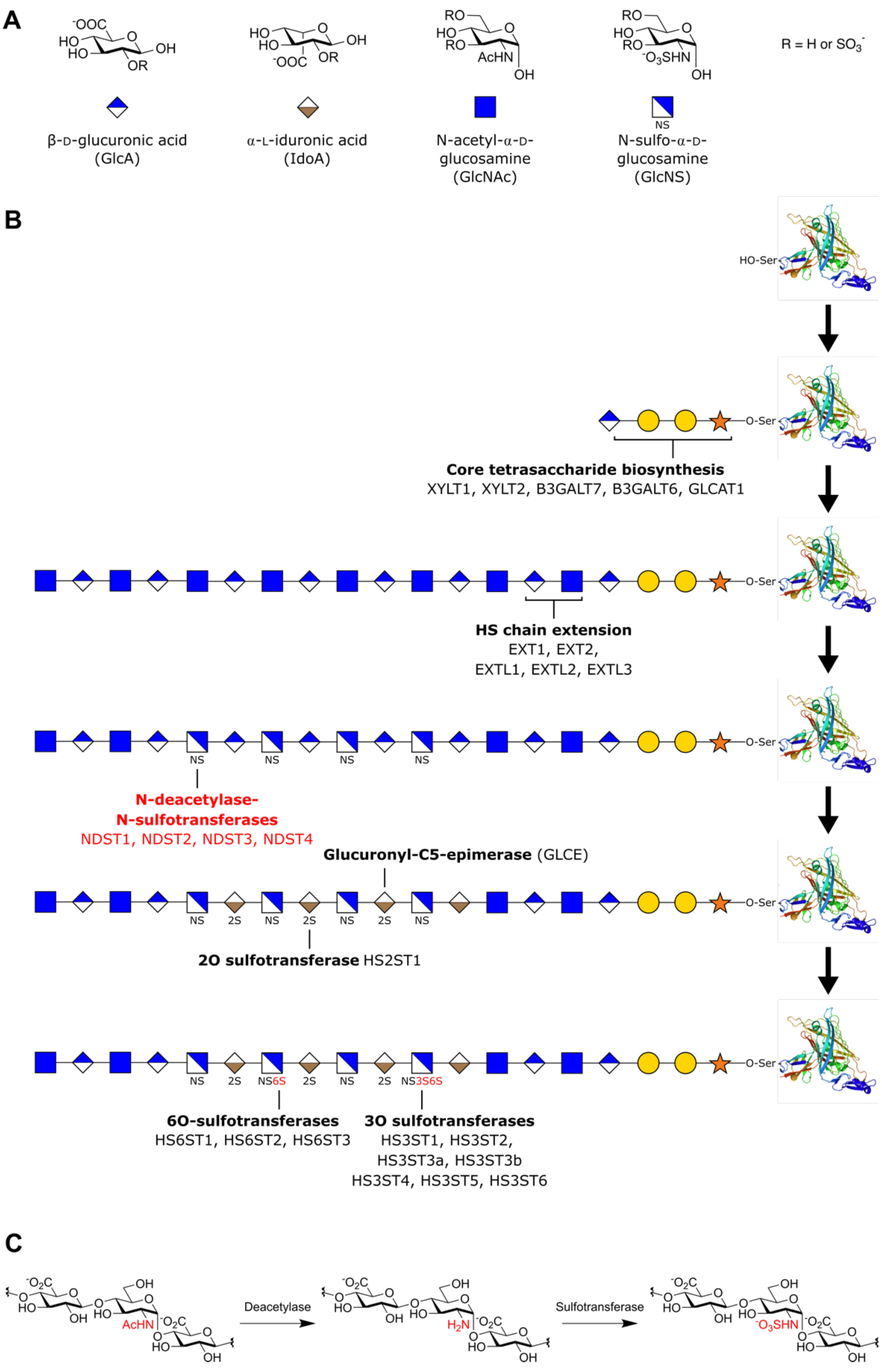


**Figure S1** Chemical structure and biosynthesis pathway of heparan sulfate (HS). (A) Monosaccharide building blocks involved in HS construction. Sites of variable O-sulfation are annotated. (B) Canonical biosynthesis pathway of HS within the Golgi complex. NDST enzymes act after chain polymerization by the EXT enzymes. (C) Bifunctionality of NDST enzymes – GlcNAc sugars are first deacetylated to glucosamine by catalytic deacetylase activity, before N-sulfation by catalytic sulfotransferase activity.

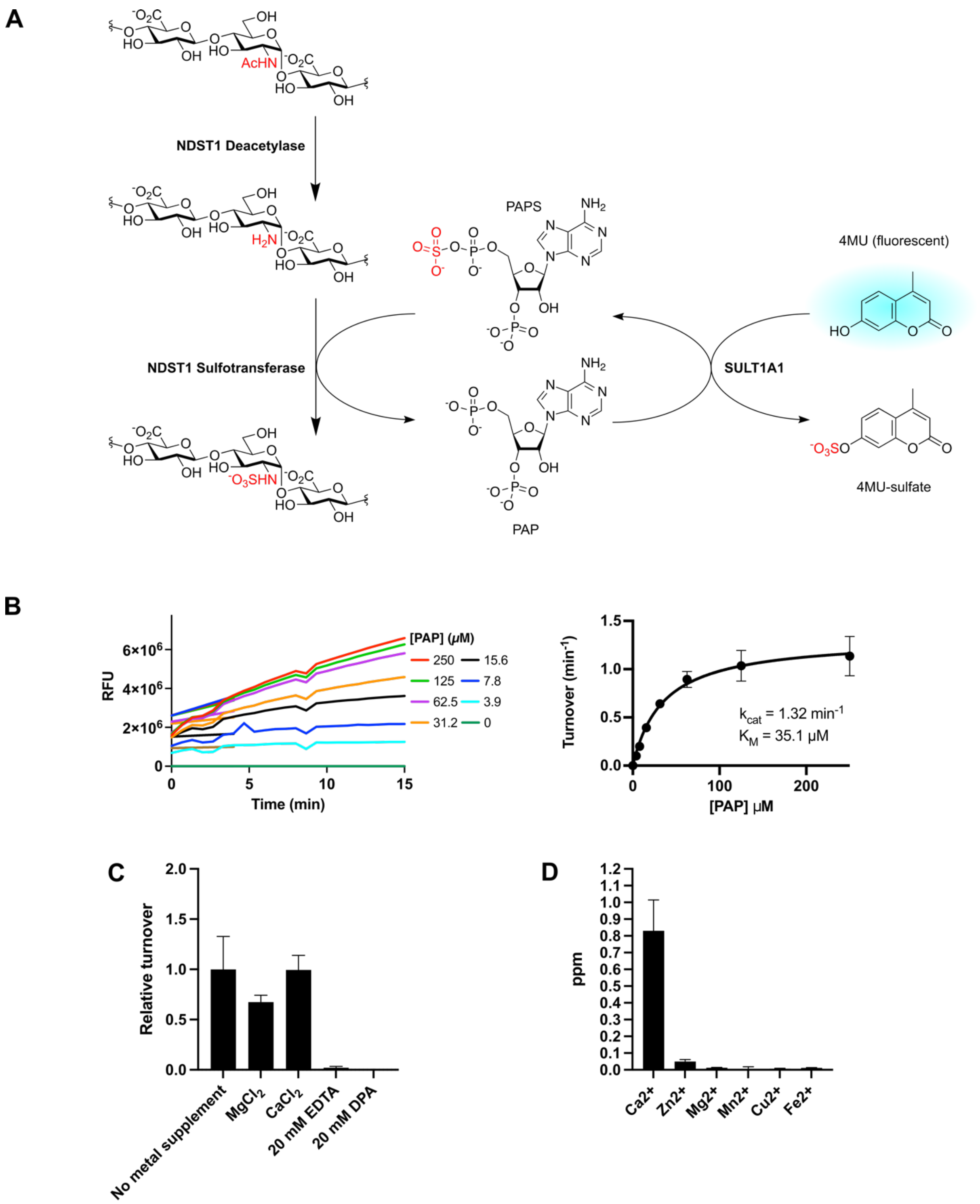


**Figure S2** Coupled enzyme assay to measure NDST1 activity. (A) Schematic of assay: 5'-phosphoadenosine-3'-phosphate (PAP) generated by NDST1 is regenerated by the bacterial enzyme Sult1A1, using fluorogenic sulfate donor 4-methylumbelliferyl (4MU) sulfate. (B) Time course measurement of Sult1A1 mediated 4MU release with respect to PAP concentration (left), Michaelis-Menten kinetics of Sult1A1 activity with respect to PAP (right). (C) Relative activity of NDST1 in the presence of $Mg^{2+}$, $Ca^{2+}$ and chelators, as measured using the Sult1A1 coupled enzyme assay. (D) ICP-OES of purified NDST1, showing the presence of carried through $Ca^{2+}$ and $Zn^{2+}$. For all bar graphs, data show mean ± standard deviation from 3 or 4 technical replicates.

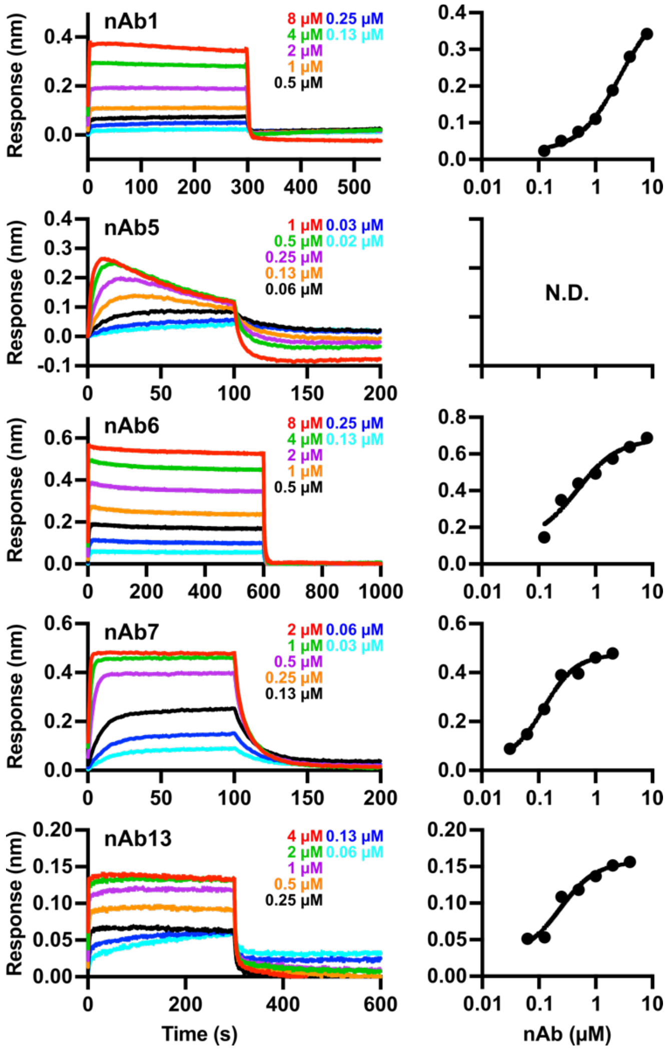


**Figure S3** Biophysical characterization of anti-NDST1 nAb affinities by biolayer interferometry (BLI). Representative sensorgrams and steady state binding curves showing interaction of nAbs with bound NDST1. Quantitated binding parameters are shown in main text **Table 1**. N.D. not determinable due to poor curve fit.

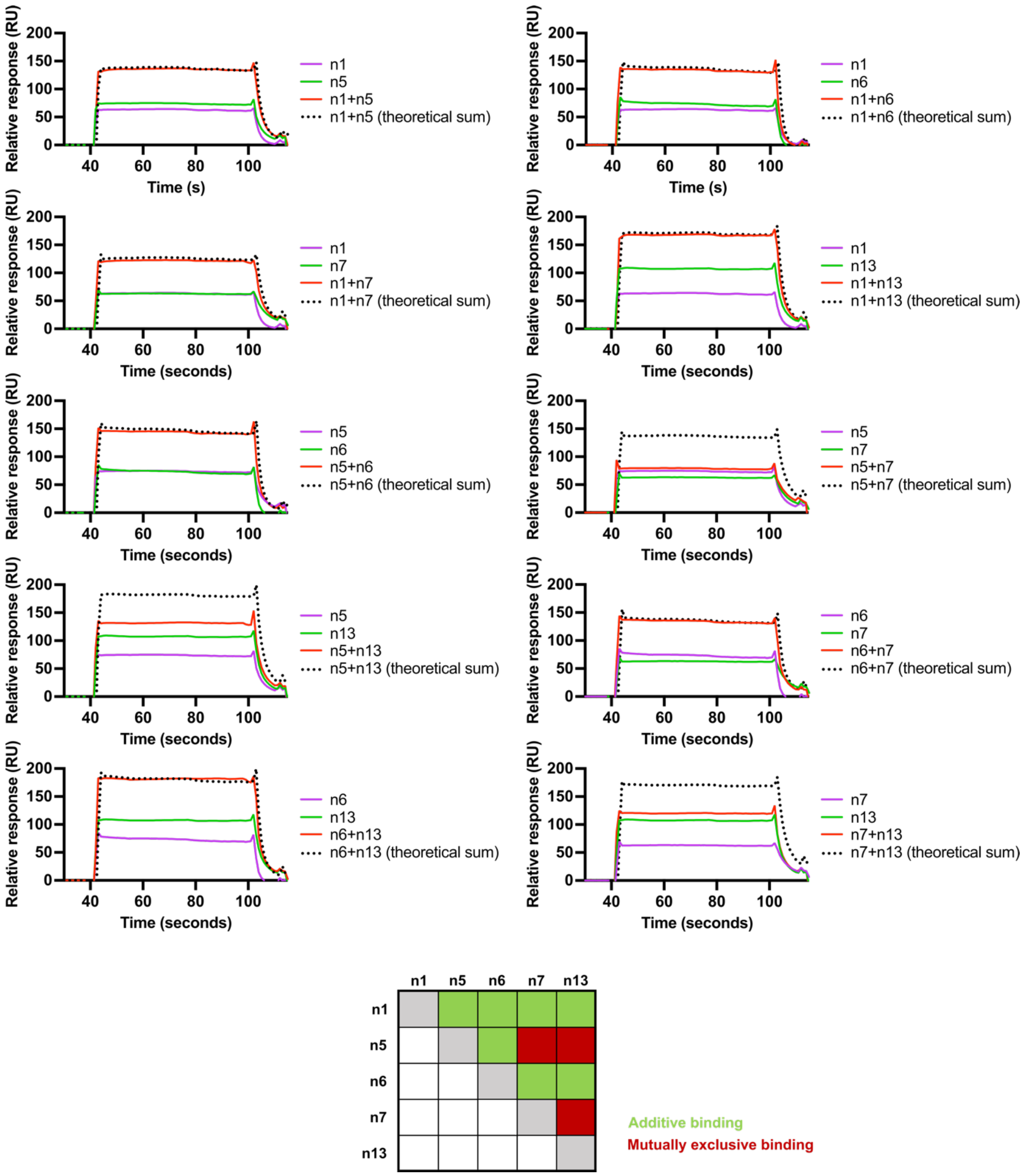


**Figure S4** Additive binding of anti-NDST1 nAbs, as measured by SPR of combinatorial nAb pairs. Binding of nAb5+nAb7, nAb5+nAb13 or nAb7+nAb13 is mutually exclusive.

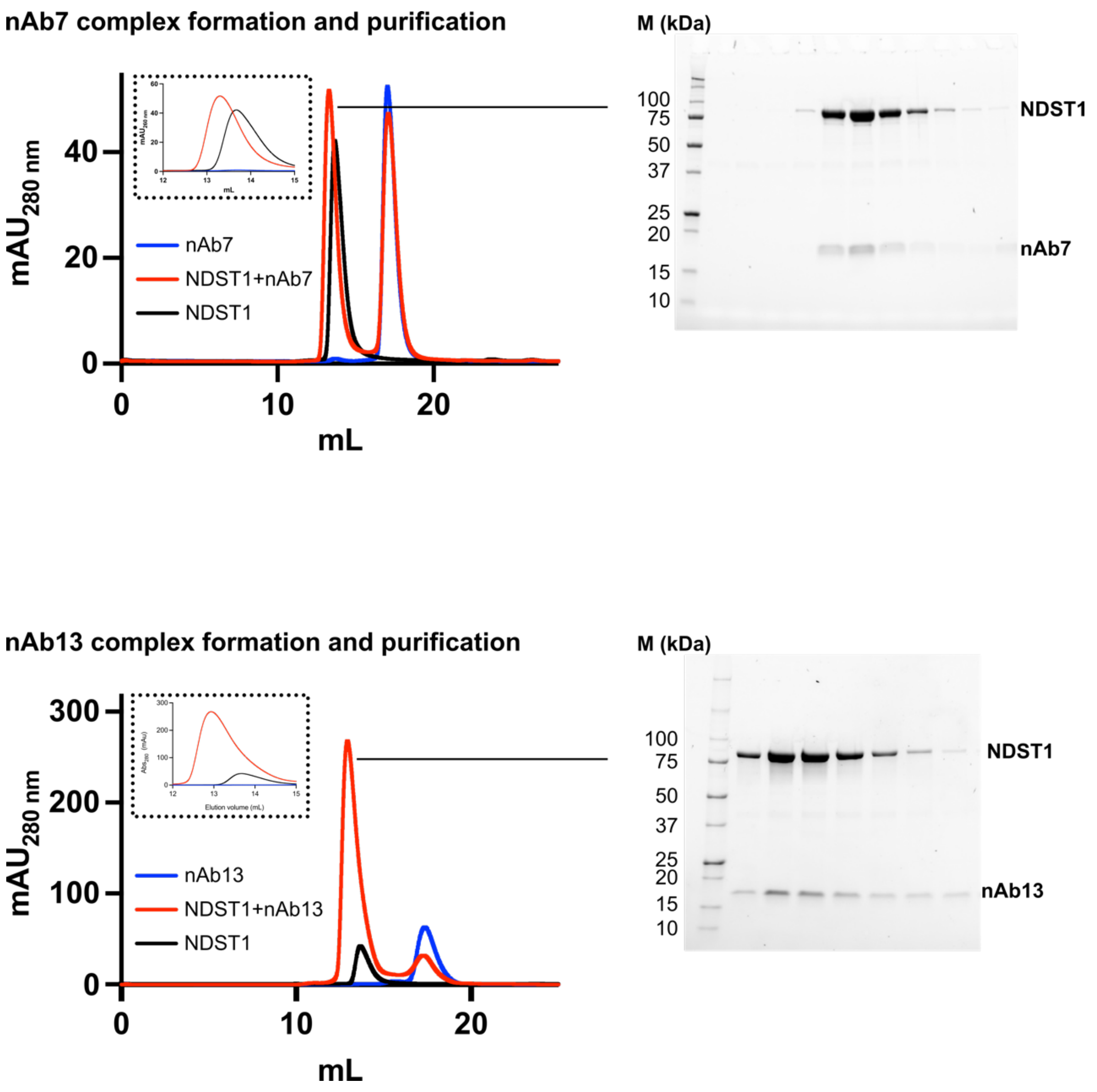


**Figure S5** SEC purification of NDST1 in complex with nAbs. nAb7 or nAb13 binding induces a shift in NDST1 retention time, indicative of stable complex formation. SDS-PAGE gel confirms co-elution of NDST1 with nAbs.

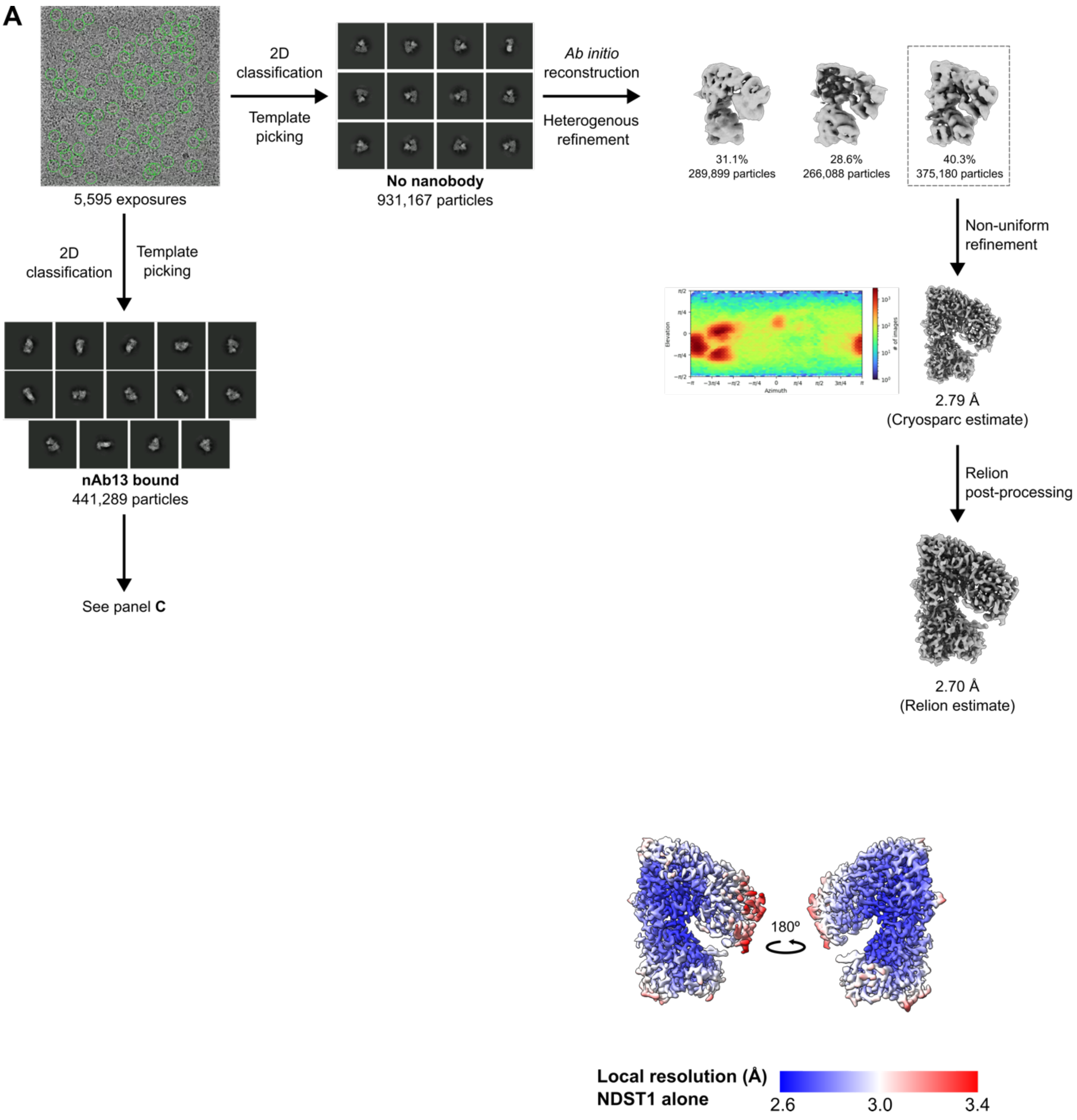
A
5,595 exposures
2D classification
Template picking
No nanobody
931,167 particles
Ab initio reconstruction
Heterogenous refinement
31.1%
289,899 particles
28.6%
266,088 particles
40.3%
375,180 particles
Non-uniform refinement
2.79 Å
(Cryosparc estimate)
Relion post-processing
2.70 Å
(Relion estimate)
2D classification
Template picking
nAb13 bound
441,289 particles
See panel C
180º
Local resolution (Å)
NDST1 alone
2.6
3.0
3.4

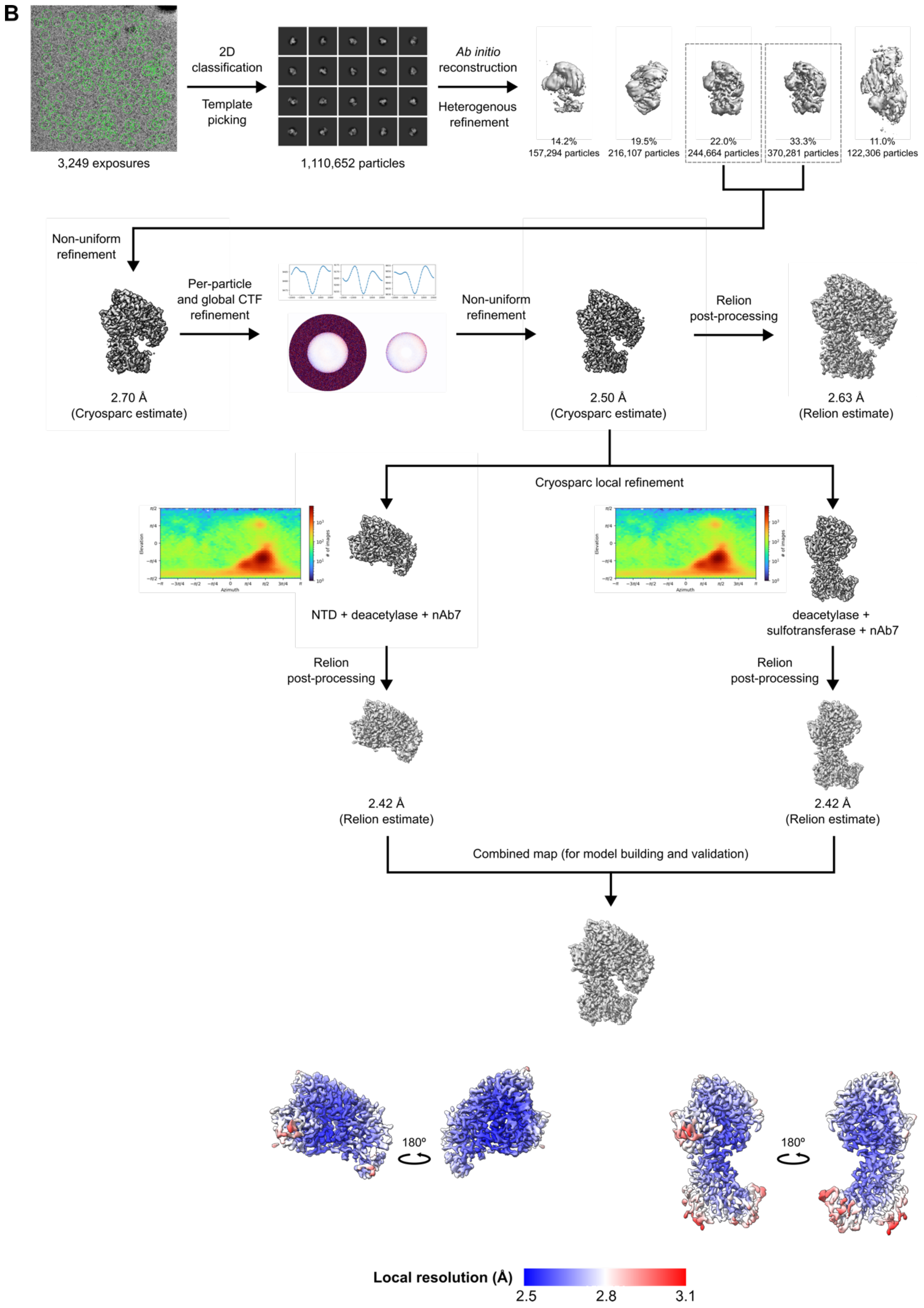

B
3,249 exposures
2D classification
Template picking
1,110,652 particles
Ab initio reconstruction
Heterogenous refinement
14.2%
157,294 particles
19.5%
216,107 particles
22.0%
244,664 particles
33.3%
370,281 particles
11.0%
122,306 particles
Non-uniform refinement
2.70 Å
(Cryosparc estimate)
Per-particle and global CTF refinement
Non-uniform refinement
2.50 Å
(Cryosparc estimate)
Relion post-processing
2.63 Å
(Relion estimate)
Cryosparc local refinement
NTD + deacetylase + nAb7
deacetylase + sulfotransferase + nAb7
Relion post-processing
2.42 Å
(Relion estimate)
Relion post-processing
2.42 Å
(Relion estimate)
Combined map (for model building and validation)
180º
180º
Local resolution (Å)
2.5
2.8
3.1

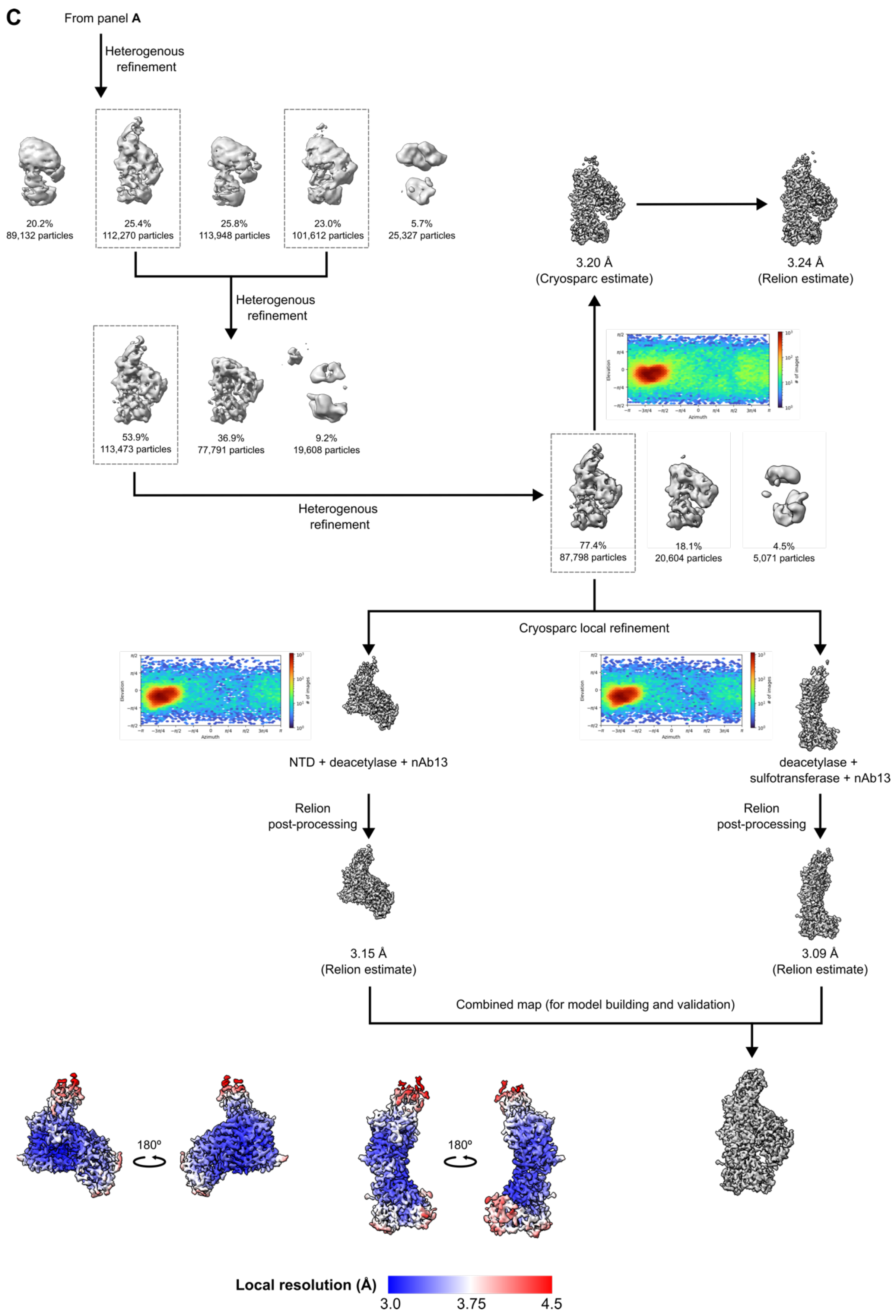
C
From panel A
Heterogenous refinement
20.2%
89,132 particles
25.4%
112,270 particles
25.8%
113,948 particles
23.0%
101,612 particles
5.7%
25,327 particles
Heterogenous refinement
53.9%
113,473 particles
36.9%
77,791 particles
9.2%
19,608 particles
Heterogenous refinement
77.4%
87,798 particles
18.1%
20,604 particles
4.5%
5,071 particles
3.20 Å
(Cryosparc estimate)
3.24 Å
(Relion estimate)
Cryosparc local refinement
NTD + deacetylase + nAb13
deacetylase + sulfotransferase + nAb13
Relion post-processing
Relion post-processing
3.15 Å
(Relion estimate)
3.09 Å
(Relion estimate)
Combined map (for model building and validation)
180º
180º
Local resolution (Å)
3.0
3.75
4.5

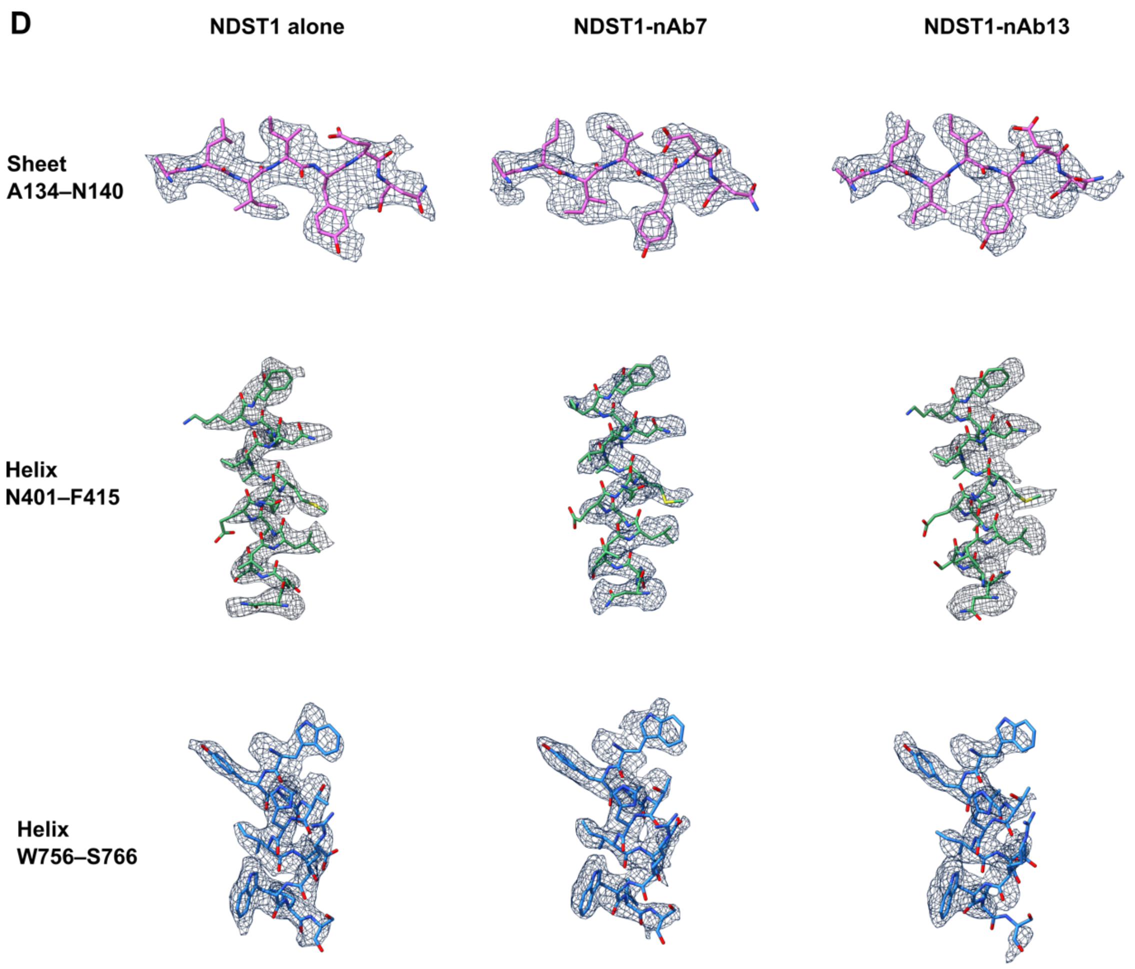


**Figure S6** Cryo-EM processing strategies and data quality. (A) nAb free NDST1. (B) NDST1-nAb7 complex. (C) NDST1-nAb13 complex. (Note that nAb free NDST1 and NDST1-nAb13 volumes were reconstructed from particles on the same grid). (C) Representative densities and model fits for structures. Maps are contoured to ChimeraX 0.12 for A134–N140, ChimeraX 0.18–0.24 for N401–F415, ChimeraX 0.14–0.15 for W756–S766.

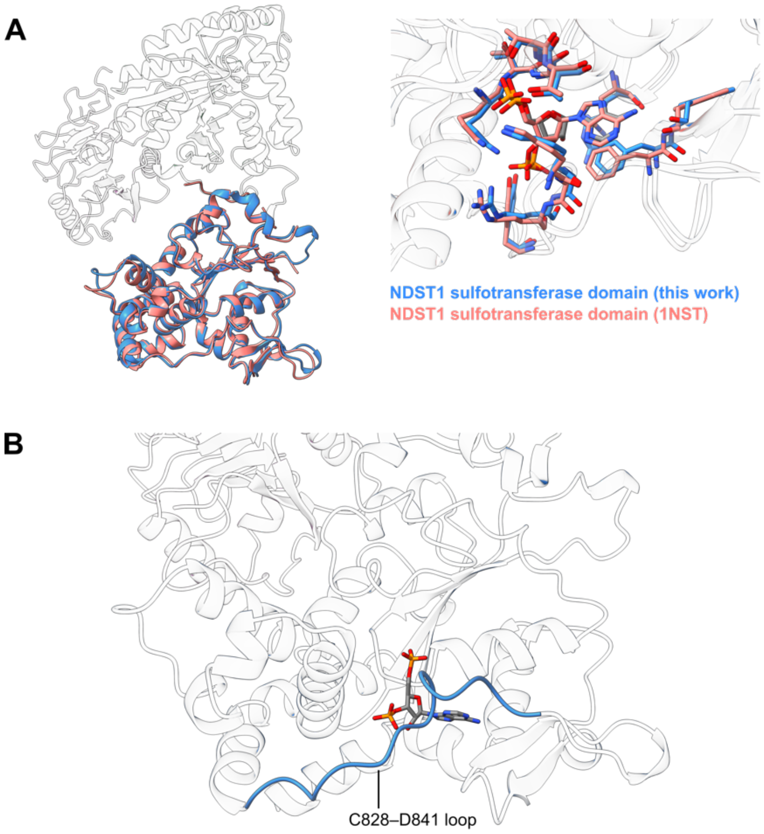


**Figure S7** Additional views of NDST1 sulfotransferase domain. (A) Comparison of NDST1 sulfotransferase domain overall fold and active site architecture with previously solved crystal structure 1NST. RMSD 0.96 Å over 277 Cαs. (B) Location of the C828–D841 loop, which appears function as a lid that holds PAP(S) within the sulfotransferase active site.

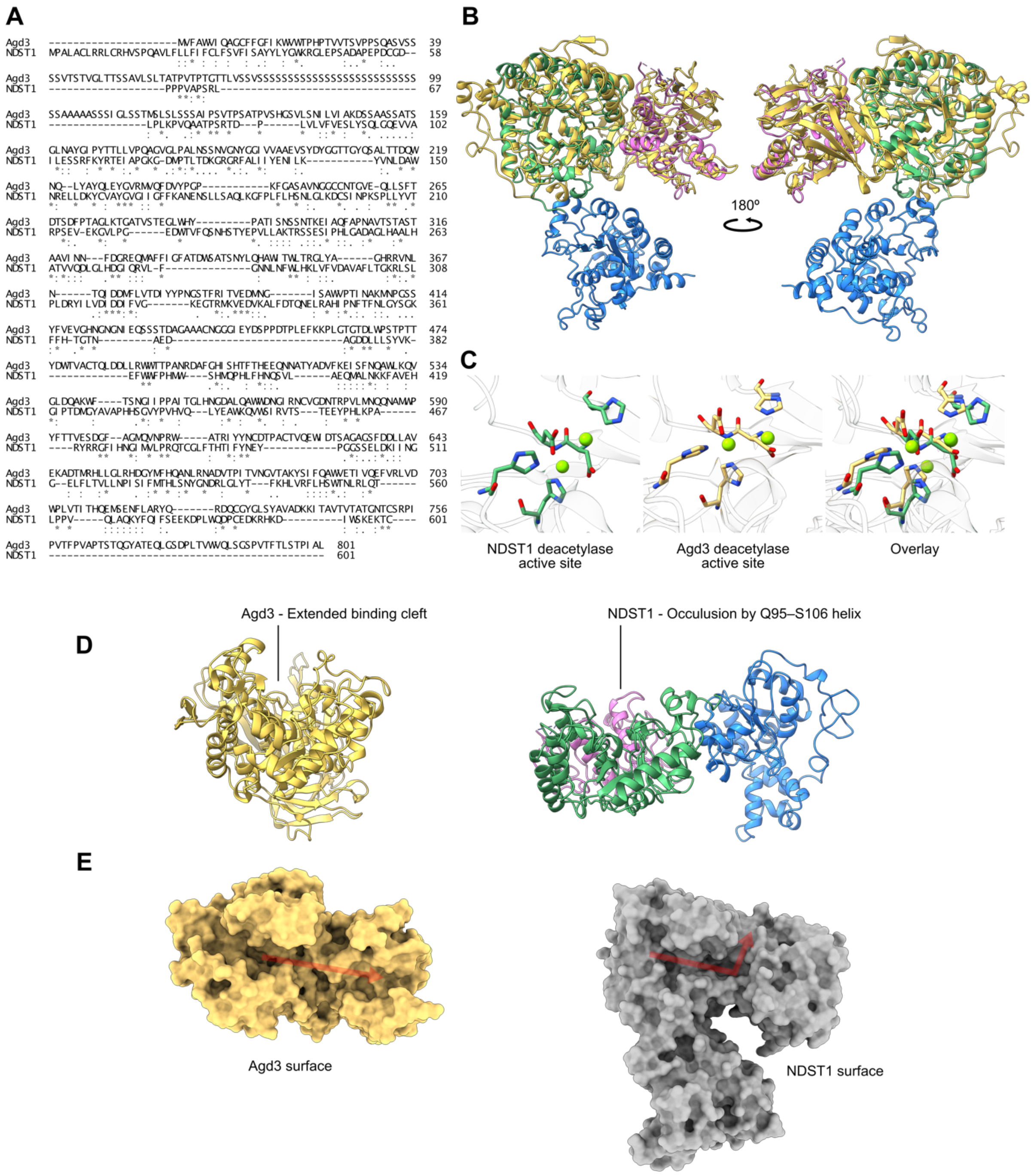


**Figure S8** NDST1 homology with Agd3 from *Aspergillus fumigatus* (PDB accession 6NWZ). (A) ClustalΩ[1] alignment of Agd3 with NDST1 residues 1-601 (N-terminal and deacetylase domains) showing 21.5% sequence identity. (B) Superposition of Agd3 and NDST1, showing similarity over N-terminal and deacetylase domains. NDST1 colored as in **Main Text Figure 3**, Agd3 colored in yellow. (C) Deacetylase domain active site comparisons for NDST1 and Agd3. Two divalent metal ions are observed close to the catalytic amino acids for both enzymes. (D)(E) Agd3 binds substrates using an extended cleft, spanning its deacetylase domain and carbohydrate binding accessory domain. A similar extended cleft in NDST1 is occluded by the NTD Q95-S106 helix. However, a binding channel can be observed at the interface between the deacetylase and N-terminal domains.

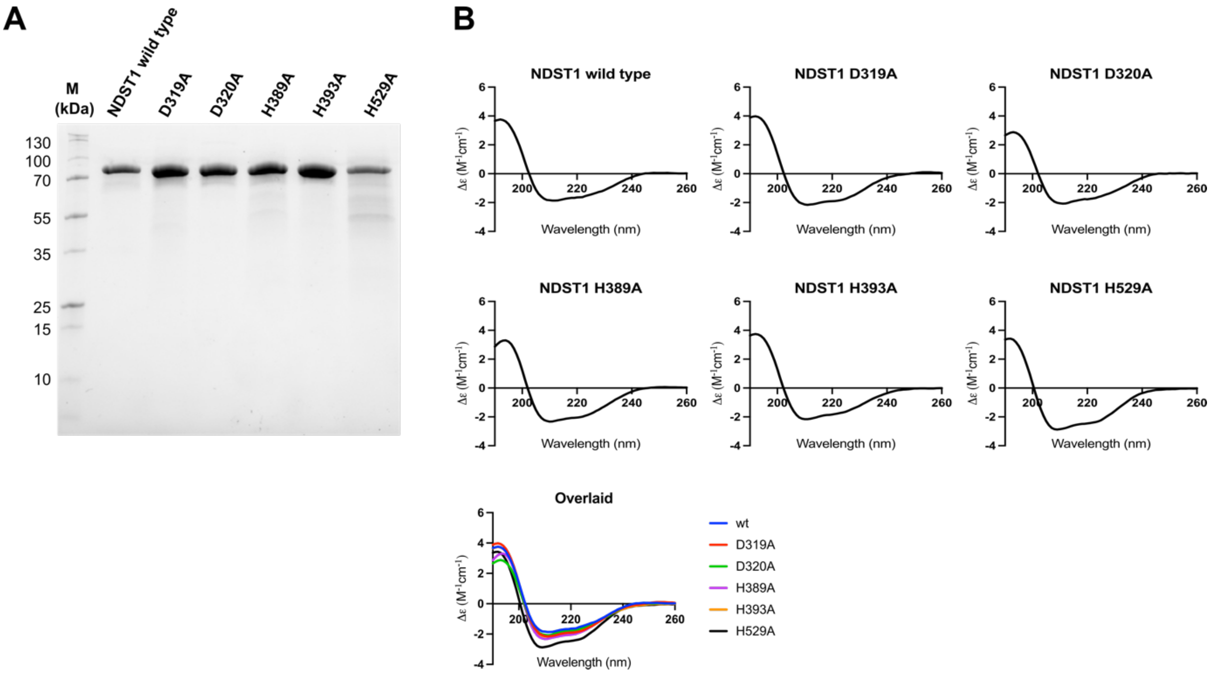


**Figure S9** SDS-PAGE (A) and circular dichroism spectra (B) indicate similar molecular mass and secondary structure composition for wild-type NDST1 vs deacetylase domain mutants.

A

B

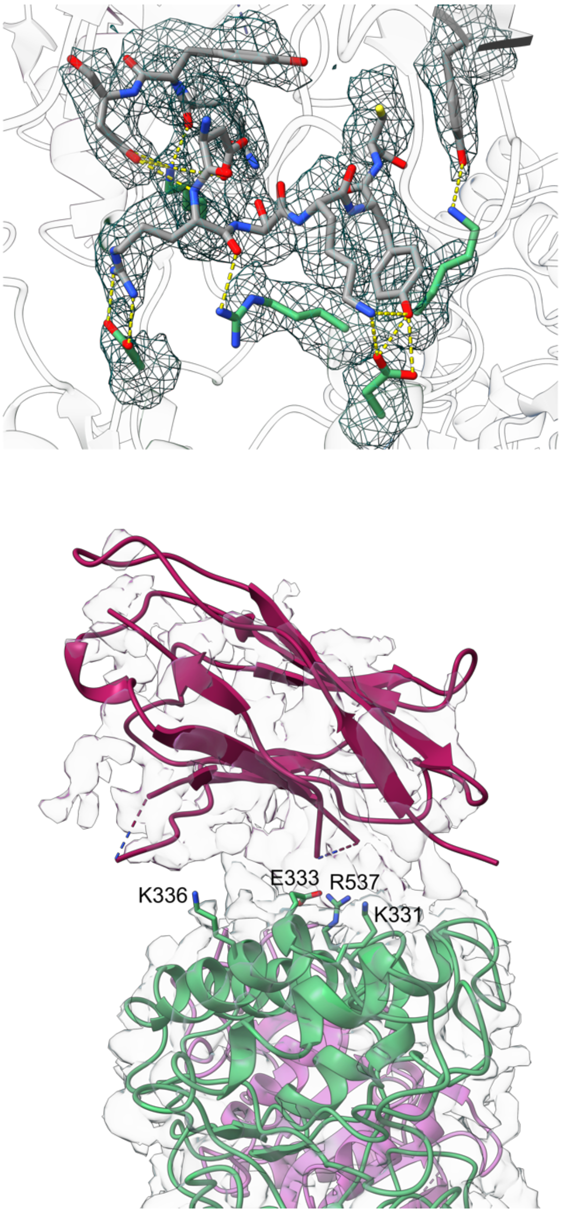


**Figure S10** cryo-EM maps of NDST1 nAb binding interfaces. (A) NDST1-nAb7 interface, with coulombic density contoured to 6.10 σ. View is identical to that in main text **Figure 3c**. (B) NDST1-nAb13 interface. NDST1 residues within 6 Å of the rigid body fitted nAb13 (K331, E333, K336 and R537) are annotated.

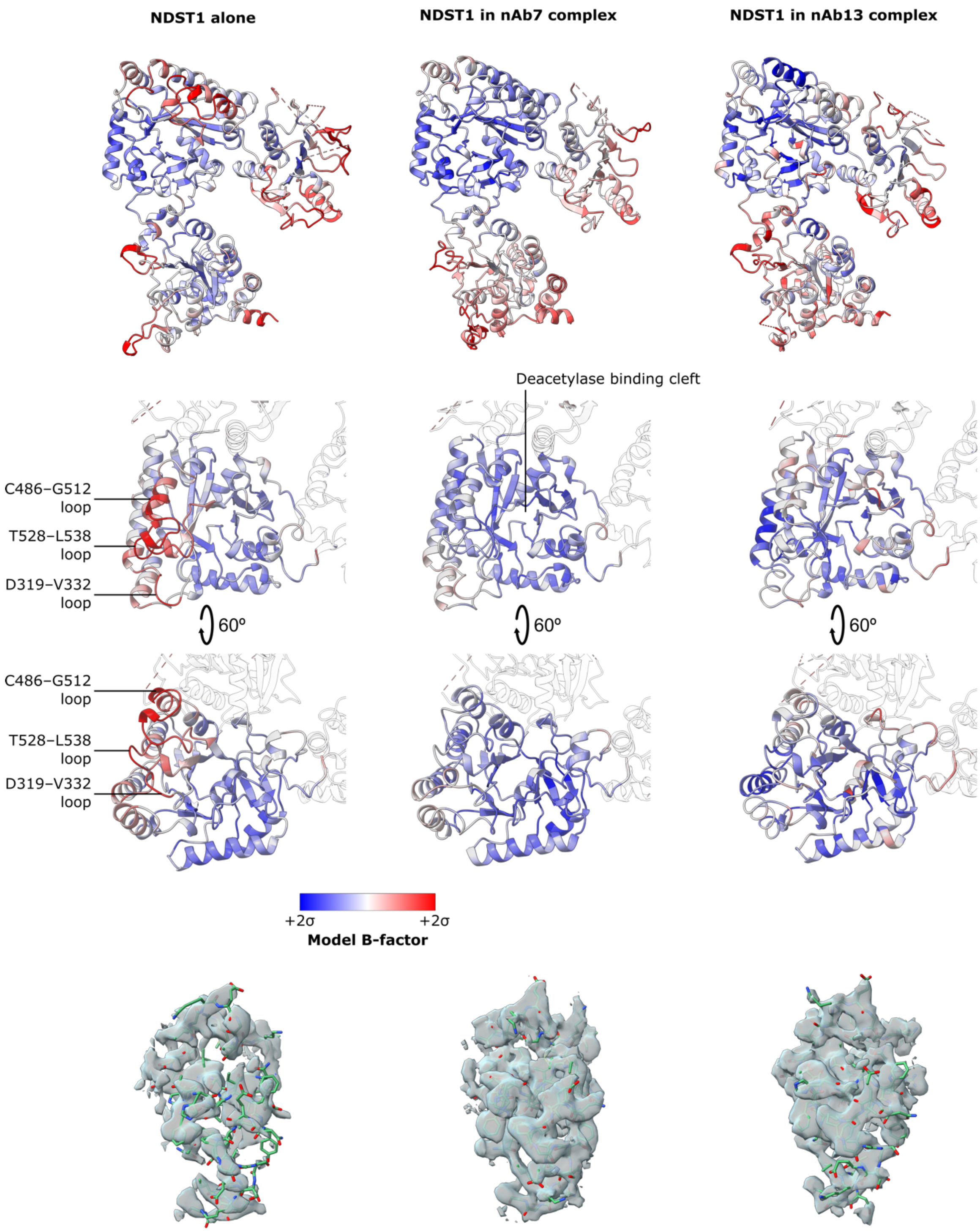


**Figure S11** B-factor analysis of NDST1 models from the NDST1 alone, NDST1-nAb7 and NDST1-nAb13 datasets. Higher B-factors (red) indicate increased structural disorder and molecular movement. Both nAb7 and nAb13 induce ordering of the D319–V332, C486–G512 and T528–L538 loops, which lie close to the deacetylase cleft, and are likely to contact substrate. Comparison of map volume around the D319–V332, C486–G512 and T528–L538 region shows nAb free NDST1 to have weaker density, indicating poor ordering of these loops. All model B factors are normalized to mean ± 2 standard deviations. Maps contoured to 0.12 in ChimeraX for all structures.

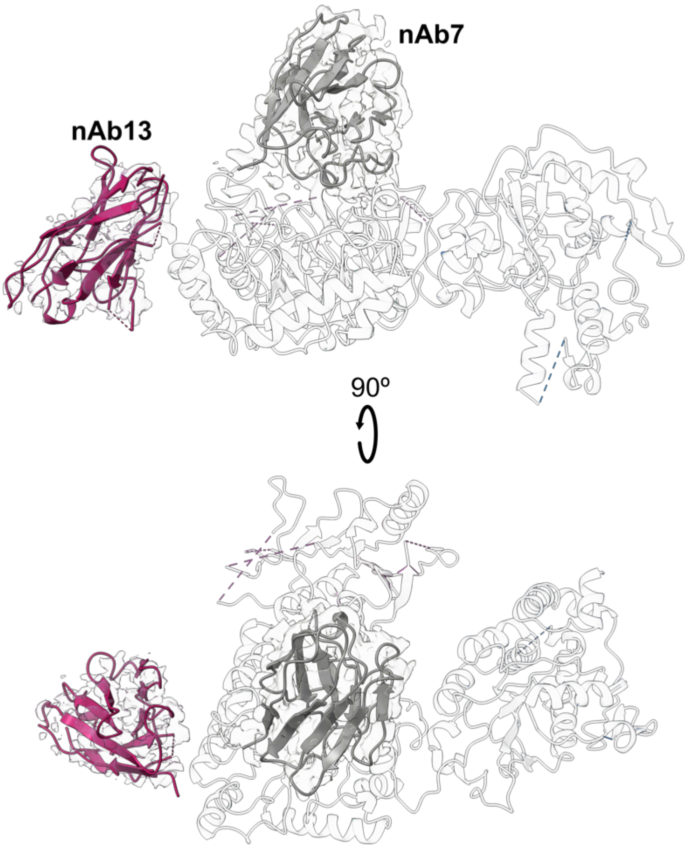


**Figure S12** Overlay of nAb7 and nAb13 binding sites on NDST1. No direct steric interference is observed between the two nAbs, suggesting their inability to mutually bind NDST1 (**Figure S4**) arises from allosteric effects.

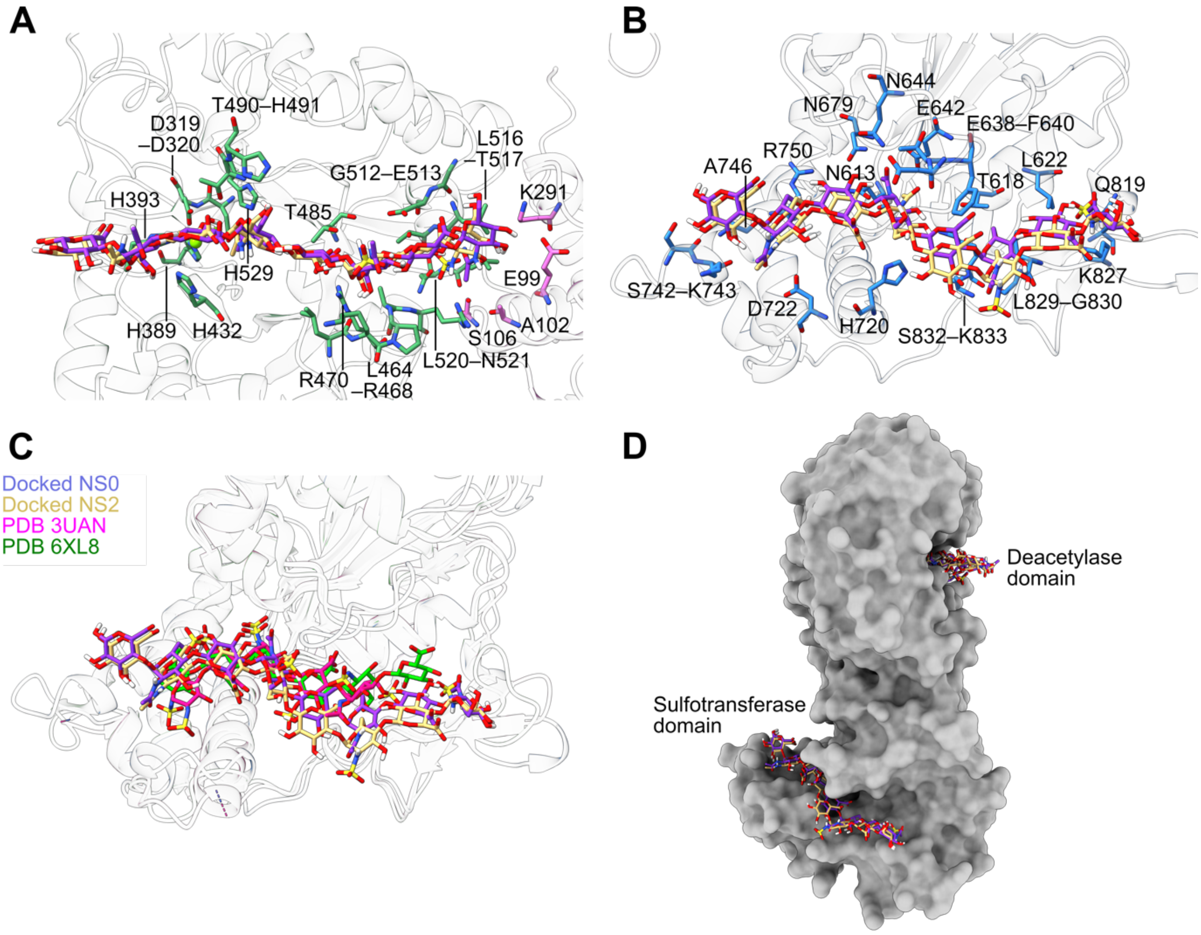


**Figure S13** Close up views of oligosaccharides docked into NDST1 active sites. (A) NS0 (purple) and NS2 (yellow) in the NDST1 deacetylase cleft, with all amino acids within 4 Å annotated. Residues from both deacetylase and N-terminal domains are present at the interaction interface. (B) NS0 and NS2 in the NDST1 sulfotransferase cleft, with amino acids within 4 Å distance annotated. (C) Comparison of docked NS0 and NS2 poses in the NDST1 sulfotransferase cleft with structures of homologous sulfotransferases HS3ST1 (3UAN) and HS3ST3 (6XL8). Trajectories of the NDST1 docked octasaccharides are similar to those experimentally observed for HS3ST1 and HS3ST3. (D) Surface overview showing the spatial relationship between oligosaccharides in the NDST1 deacetylase and sulfotransferase domains.

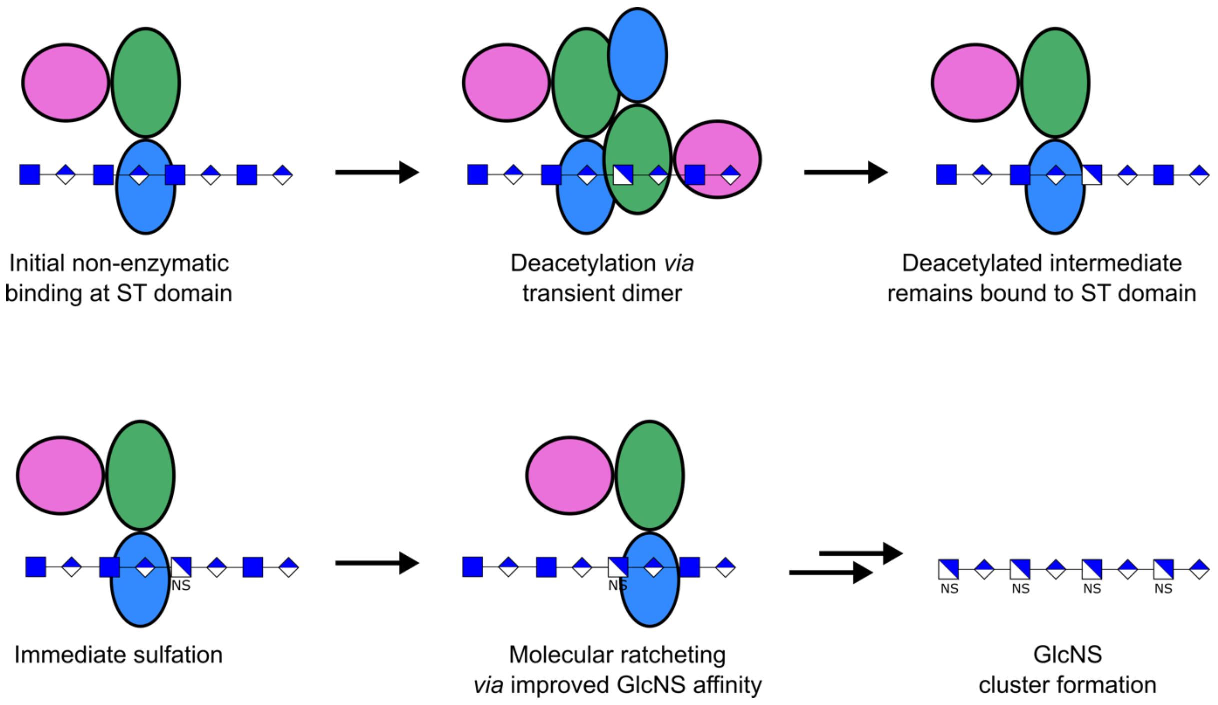


**Figure S14** Putative model for bifunctional NDST1 catalysis. Initial HS binding at the NDST1 sulfotransferase domain occurs prior to enzymatic processing. Deacetylation by transient dimerization leads to the formation of a GlcN intermediate, which is immediately processed by the vicinal sulfotransferase active site. Because sulfotransferase domain interactions are electrostatic, they may interact more strongly with GlcNS residues, producing a ratcheting effect that can drive GlcNS cluster formation. Domain colors correspond to those in **Main Text Figure 3**.

**Table S1** Cryo-EM data processing and model building statistics

| | NDST1 alone<br><br>(EMD-16564)<br>(PDB 8CCY) | NDST1-nAb7<br>NTD–DeAc–nAb7 locally refined map<br>(EMD-16627) | NDST1-nAb7<br>DeAc–ST–nAb7 locally refined map<br>(EMD-16629) | NDST1-nAb7<br>Composite map for model building<br>(EMD-16565)<br>(PDB 8CD0) | NDST1-nAb13<br>NTD–DeAc–nAb13 locally refined map<br>(EMD-16662) | NDST1-nAb13<br>DeAc–ST–nAb13 locally refined map<br>(EMD-16663) | NDST1-nAb13<br>Composite map for model building<br>(EMD-16664)<br>(PDB 8CHS) |
|---|---|---|---|---|---|---|---|
| **Data collection and processing** | | | | | | | |
| Magnification | 165,000x | 165,000x | 165,000x | | 165,000x | 165,000x | |
| Voltage (kV) | 300 | 300 | 300 | | 300 | 300 | |
| Electron exposure (e–/Å$^2$) | 50 | 50 | 50 | | 50 | 50 | |
| Defocus range (μm) | -1.2 to -2.6 | -1.2 to -2.6 | -1.2 to -2.6 | | -1.2 to -2.6 | -1.2 to -2.6 | |
| Pixel size (Å) | 0.73 | 0.73 | 0.73 | | 0.73 | 0.73 | |
| Symmetry imposed | C1 | C1 | C1 | | C1 | C1 | |
| Initial particle images (no.) | 3,867,804 | 2,721,982 | 2,721,982 | | 3,867,804 | 3,867,804 | |
| Final particle images (no.) | 371,778 | 614,944 | 614,944 | | 87,798 | 87,798 | |
| Map resolution (Å) | 2.70 | 2.42 | 2.42 | | 3.15 | 3.09 | |
| FSC threshold | 0.143 | 0.143 | 0.143 | | 0.143 | 0.143 | |
| Map resolution range (Å) | 2.66–4.58 | 2.53–3.45 | 2.58–3.49 | | 3.01–5.70 | 3.14–6.27 | |
| **Refinement** | | | | | | | |
| Initial model used (PDB code) | Alphafold2 prediction | | | NDST1 alone model; 7TGF | | | NDST1 alone model; nAb7 (CA only) |
| Model resolution (Å) | 2.9 | | | 3.2 | | | 4.0 |
| FSC threshold | 0.5 | | | 0.5 | | | 0.5 |
| Map sharpening *B* factor (Å$^2$) | 0 | | | 0 | | | 0 |
| Model composition | | | | | | | |
| Non-hydrogen atoms | 6235 | | | 6860 | | | 6424 |
| Protein residues | 754 | | | 837 | | | 820 |
| Ligands | 3 | | | 2 | | | 2 |
| *B* factors (Å$^2$) | | | | | | | |
| Protein | 85.7 | | | 88.8 | | | 62.1 |
| Ligand | 74.0 | | | 133.2 | | | 77.9 |
| R.m.s. deviations | | | | | | | |
| Bond lengths (Å) | 0.005 | | | 0.007 | | | 0.008 |
| Bond angles (°) | 0.928 | | | 1.25 | | | 1.65 |
| Validation | | | | | | | |
| MolProbity score | 1.36 | | | 1.62 | | | 1.95 |
| Clashscore | 3.82 | | | 3.48 | | | 5.94 |
| Poor rotamers (%) | 1.35 | | | 2.03 | | | 2.19 |
| Ramachandran plot | | | | | | | |
| Favored (%) | 97.58 | | | 96.34 | | | 94.64 |
| Allowed (%) | 2.42 | | | 3.66 | | | 5.36 |
| Disallowed (%) | 0.00 | | | 0.00 | | | 0.00 |

### Methods

*Cloning and gene expression – NDST1*

A plasmid containing cDNA encoding the NDST1 gene was obtained from the DNASU repository (Arizona State University), from which a cDNA fragment encoding solubilized NDST1 (residues 79–882) was amplified using Q5 high fidelity DNA polymerase (New England Biolabs). The NDST1(79–882) fragment was subcloned using Infusion (Takara Bio) into the pOMNIBac transfer vector (Geneva Biotech), behind a honeybee melittin secretion peptide, 6xHis tag and TEV cleavage site. NDST1 mutants were derived from the wild-type NDST1(79–882) plasmid by site-directed mutagenesis using a PCR based method[2]. cDNA encoding for C-terminal Avi-tagged NDST1 for biotinylation was generated by PCR with tailed primers, followed by subcloning into pOMNIBac. All constructs were verified by Sanger sequencing before further use.

Recombinant bacmids were produced *via* the Tn7 transposition method in DH10EMBacY cells (Geneva Biotech), and purified using the PureLink miniprep kit (Invitrogen) according to manufacturers protocols. V1 baculovirus stocks were produced by transfection of bacmid into low-passage ExpiSf9 cells (Invitrogen) in adherent format using FuGENE HD (Promega) at a ratio of ~2 mg bacmid DNA to ~5 µL FuGENE. V1 to V2 virus amplification was carried out in ExpiSf9 cells in suspension culture, at a density of ~1-2 x $10^6$ $mL^{-1}$. Viral propagation was tracked using the YFP marker encoded by the EMBacY virus. Optimum baculovirus amplification was typically achieved 72 h after inoculation, when cells appeared 60+% fluorescent. V2 virus stocks were harvested by centrifugation at 200 g for 5 min at 4°C to remove cells, and supplemented with 2% v/v FBS for storage.

Large scale gene-expressions were carried out in High-Five (Invitrogen; Trichoplusia ni) or ExpiSf9 cells in log-phase growth. V2 baculovirus was added to cells at a multiplicity of infection > 1, and the resulting infection tracked using the EMBacY YFP marker. Cultures were typically harvested at ~72 h, or when cells showed ~80% or greater fluorescence.

*Protein purification – NDST1*

Solubilized recombinant NDST1 was purified from conditioned insect cell media. Typically, ~2–3 L of insect cell culture was cleared of cells by centrifugation at 200 x g for 15 min at 4°C, followed by further clearing of debris by centrifugation at 5,000 x g for 60 min at 4°C. Clarified media was supplemented with DL-dithiothreitol (DTT; 1 mM) and phenylmethylsulfonyl fluoride (PMSF; 0.2 mM), before loading onto a 5 mL Histrap Excel column (Cytiva), pre-equilibrated with Histrap buffer A (20 mM Tris pH 8.0, 500 mM NaCl, 20 mM Imidazole, 1 mM DTT), at a rate of 5 $mL.min^{-1}$. The loaded Histrap column was washed with 10 column volumes (CV) Histrap buffer A, before eluting with a linear gradient of Histrap buffer A to Histrap buffer B (20 mM Tris pH 8.0, 500 mM NaCl, 500 mM Imidazole, 1 mM DTT) over 10 column volumes (CV). NDST1 containing fractions, as determined by SDS-PAGE, were pooled, diluted >10-fold in milli-Q water, and adjusted to pH 6.5 by the addition of 1 M MES acid, before loading onto a Hitrap SP XL column (Cytiva) pre-equilibrated with ion exchange (IEX) buffer A (20 mM MES pH 6.5, 1 mM DTT). Loaded Hitrap SP XL columns were washed with 10 column volumes (CV) IEX buffer A, before eluting with a linear gradient of IEX buffer A to IEX buffer B (20 mM MES pH 6.5, 1500 mM NaCl, 1 mM DTT) over 10 CV. NDST1 containing fractions were pooled again and digested overnight at room temperature with TEV protease at 1:100 mass ratio TEV:NDST1 to cleave the 6xHis tag. TEV digested NDST1 was rerun over a 5 mL Histrap Excel column pre-equilibrated with Histrap buffer A, which was then washed with 4 CV Histrap buffer A. Flowthrough and wash fractions were pooled and concentrated to 2 mL volume using a 30 kDa molecular weight cut-off (MWCO) Vivaspin concentrator (Cytiva). Final purification was carried out by size-exclusion chromatography (SEC), using a HiLoad 16/600 Superdex 200 pg column (Cytiva), pre-equilibrated and run in SEC buffer (20 mM MES pH 6.5, 200 mM NaCl, 1 mM DTT). Pure NDST1 containing fractions were pooled, adjusted to <100 mM NaCl using IEX buffer A, and concentrated to ~10 mg/mL using a 30 kDa MWCO Vivaspin concentrator. Small aliquots were flash frozen using liquid nitrogen, and stored at -80°C for further use.

Mutant NDST1 constructs were purified in the same way as for wild type protein. Biotinylated NDST1 was prepared using purified C-terminal avitagged protein, which was treated with the BirA biotin-protein ligase reaction kit (Avidity, LLC) following the manufacturers protocol. Biotinylated protein was purified by a final round of SEC following the BirA reaction.

*Cloning and gene expression – SULT1A1*

A plasmid encoding cDNA for SULT1A1 from *Rattus norvegicus* was obtained from GenScript Biotech. A cDNA fragment encoding for wild type SULT1A1 was subcloned into the pET15b backbone, behind a thrombin cleavage site, using Ndel and BamHI restriction sites. Mutations K65E and R68G, which prevent the formation of an inhibitory SULT1A1-PAP-4-methylumbelliferone complex[3], were introduced into the plasmid by site-directed mutagenesis. The final construct was verified by Sanger sequencing before further use.

For expression, SULT1A1-K65E-R68G was used to transform BL21 Gold (DE3) chemically competent cells (Agilent) by heat shock. Transformed cells were grown in TB media containing 100 $\mu g.mL^{-1}$ ampicillin at 37°C with shaking, until reaching an $OD_{600}$ of 0.8–1.0. Cultures were induced by the addition of isopropyl-b-thiogalactoside (IPTG; Generon) to a final concentration of 0.5 mM, then grown overnight at 16°C with shaking. Cultures were harvested by centrifugation at 5000 g for 15 min at 4°C, and pellets stored at –80°C until further processing.

*Protein purification – SULT1A1-K65E-R68G*

SULT1A1-K65E-R68G pellets were thawed and resuspended in 100 mM Tris pH 7.6, 1 mM DTT containing 120 kU DNase I (Sigma) and cOmplete™, EDTA-free Protease Inhibitor Cocktail (Sigma). Resuspended cell pellets were lysed via two rounds of cell disruption at 30 kpsi (Constant Systems) at 4°C. Cell debris was removed by centrifugation at 14,000 g at 4°C for 30 min, and supernatant filtered prior to loading onto a 5 mL Histrap FF column, pre-equilibrated in Histrap buffer A (100 mM Tris pH 7.6, 300 mM NaCl, 20 mM imidazole, 1 mM DTT). The loaded column was washed with 5 CV of buffer A prior to elution of SULT1A1-K65E-R68G using a linear gradient of 0-100% Histrap buffer B (100 mM Tris pH 7.6, 300 mM NaCl, 250 mM imidazole, 1mM DTT) over 10 CV. SULT1A1-K65E-R68G containing fractions were pooled and digested overnight at room temperature with thrombin at 1:100 mass ratio thrombin:SULT1A1-K65E-R68G. Thrombin cleaved SULT1A1-K65E-R68G was passed over a 5 mL Histrap FF column pre-equilibrated in buffer A, and the column was washed with 5 CV of the same buffer. The flow-through and wash, containing de-tagged SULT1A1-K65E-R68G, were collected and pooled before being concentrated using a 10 kDa MWCO spin filter. SULT1A1-K65E-R68G was further purified by SEC using a HiLoad 16/600 Superex 75 pg column pre-equilibrated in SEC buffer (100 mM Tris pH 7.6, 1mM DTT). Fractions containing SULT1A1-K65E-R68G were assessed by SDS-PAGE and Coomassie staining before being pooled and concentrated to 6 mg.mL$^{-1}$ using a 10 kDa MWCO spin filter. Aliquots of concentrated SULT1A1-K65E-R68G were flash frozen in $LN_2$ and stored at -80°C until further use.

*Expression – heparin lyase II*

The gene encoding heparin lyase II from *Pedobacterium heparinus,* inserted into pRSET A (gifted from Marcelo Lima, University of Keele), was transformed into chemically competent C41(DE3) pLysS cells by heat shock. Pre-cultures of transformed cells were grown in LB media containing 100 µg.mL$^{-1}$ ampicillin and 50 µg.mL$^{-1}$ chloramphenicol for 16 hours, 37°C, 250 rpm. 1 L cultures of TB media, containing 100 µg.mL$^{-1}$ ampicillin and 50 µg.mL$^{-1}$ chloramphenicol were inoculated with the pre-culture and grown at 37°C, 250 rpm with shaking until reaching an $OD_{600}$ of 0.6–0.8. Cultures were cooled to 22°C and induced with 0.5 mM IPTG before further incubation at the same temperature for 16 hours, 250 rpm with shaking. Cultures were harvested at 6000 g for 15 minutes at 4°C. Pellets were stored at –80°C until further use.

*Protein purification – heparin lyase II*

Thawed cell pellets were resuspended in 20 mM sodium phosphate pH 7.9, 500 mM NaCl, 5 mM imidazole supplemented with 120 kU DNAse and cOmplete EDTA-free protease inhibitor. Cells were lysed via two rounds of cell disruption at 30 kpsi, then centrifuged at 14,000 g, 4°C, for 30 minutes to remove cell debris. The supernatant was filtered and loaded onto a 5 mL HisTrap FF column pre-equilibrated in Histrap buffer A (20 mM sodium phosphate pH 7.9, 500 mM NaCl, 5 mM imidazole), which was then washed with 5 CV of the same buffer prior to elution with a linear gradient of 0-100% Histrap buffer B (20 mM sodium phosphate pH 7.9, 500mM NaCl, 250 mM imidazole) over 10 CV. Fractions containing heparinase lyase IIwere pooled, diluted with 10 volumes of IEX buffer A (20 mM MES pH 6.0) and then loaded onto a 5 mL Hitrap SP XL column (Cytiva) pre-equilibrated in the same buffer. Elution was conducted with a linear gradient of 0-100% IEX buffer B (20 mM MES pH 6.0, 1.5 M NaCl) over 20 CV. Fractions containing heparin lyase II were pooled and concentrated using a 30 KDa MWCO spin filter, then allied to a HiLoad 16/600 Superex 200 pg column equilibrated and run with SEC buffer (20 mM sodium phosphate pH 6.8, 150 mM NaCl). Eluted heparin lyase II containing fractions were assessed by SDS-PAGE, before being pooled and concentrated to ~2 mg/mL using a 30 KDa MWCO spin filter. Heparin lyase II was stored at 4°C until required.

*Circular Dichroism*

The CD spectrum of 250 $\mu g.mL^{-1}$ NDST-1, and mutants thereof, were acquired in 20 mM phosphate buffer pH 7.4 using a Chirascan V100 (Applied Photophysics), equipped with a 0.2-mm path length quartz cuvette (Hellma, United States). All spectra were recorded with a scanning speed of 1 $nm.s^{-1}$ with 0.5 nm resolution, between 190 and 260nm using Applied Photophysics System 2.02 software. Spectra are presented as the mean of five independent scans smoothed with second-order polynomial smoothing through 11 neighbours, using GraphPad Prism 9.

*Purification of K5 polysaccharide*

K5 polysaccharide was purified from *E. coli* strain Bi 8337/41(O10:K5:H4). Typically, 1 L of LB media was inoculated with 10 mL of *E. coli* strain Bi 8337/41(O10:K5:H4), grown for 16 hours at 37°C with shaking, before being incubated for a further 55 hours at 37°C with shaking. Cells were removed via centrifugation at 8000 g for 15 min at 4°C and the supernatant collected. The supernatant was filtered and diluted with 1 volume of IEX buffer A (20 mM sodium acetate pH 4.0, 50 mM NaCl), further adjusted to pH 4.0 with acetic acid as required, before being loaded onto a C 10/20 column (Cytiva) packed with DEAE Sepharose resin (Biorad), pre-equilibrated with the same buffer. Following loading, the column was washed with IEX buffer A and bound K5 polysaccharide eluted in a single step with IEX buffer B (20 mM sodium acetate pH 4.0, 1 M NaCl). The eluant was mixed with 3 volumes of ice-cold absolute ethanol and stored overnight at –20°C. The resulting precipitate containing K5 polysaccharide was collected by centrifugation at 13,000 g, 4°C for 30 minutes and resuspended in $dH_2O$ before being lyophilised. Lyophilized K5 was dissolved in 1 M NaCl to 15 $mg.mL^{-1}$ and pH adjusted to 9.5 with 1 M NaOH. 30% hydrogen peroxide was added to a final concentration of 1.5%, before incubation overnight at room temperature. Bleached K5 polysaccharide was recovered by the addition of 3 volumes of ice-cold ethanol followed by storage at -20°C overnight. The precipitate was collected by centrifugation at 13,000 g at 4°C for 30 minutes and dialysed against $dH_2O$, exchanged a minimum of three times, using a 5 kda MWCO dialysis membrane (Biodesign). Dialysed K5 polysaccharide was filtered and lyophilised before being stored at -20°C for further use. The purity of K5 was assessed using nuclear magnetic resonance (NMR). Polysaccharide was exchanged into $D_2O$ (700 µL) and 1-dimensional ($^1H$) spectra were recorded using a Bruker Avance NEO 600 MHz spectrometer fitted with a BBOH&F cryoprobe at 298 K.

*Fluorometric assay for determination of SULT1A1-K65E-R68G kinetics*

An 8-point 2-fold serial dilution of PAP from 250 µM was incubated with 350 nM SULT1A1-K65E-R68G, 4 mM 4-methylumbellferyl (4MU)-sulfate in 50 mM Tris-HCl pH 7.5, 15 mM $MgCl_2$, 1 mM DTT, in black half area 96 well plates (Greiner) with a total volume of 20 µL. Controls omitting PAP were included and subtracted from each reaction prior to further analysis. Reactions were initiated by the addition of SULT1A1-K65E-R68G and the change in fluorescence recorded at using a Clariostar plate reader (BMG Labtech) using the 4MU fluorescence preset (Ex. 360 [b.p. 20] nm; Em. 450 [b.p. 30] nm). An 8 point 2-fold serial dilution standard curve of 4MU (from 10 µM) in 50 mM Tris-HCl pH 7.5, 15 mM $MgCl_2$ was used to calculate the evolution of 4MU during the reaction course. Initial reaction velocities at each substrate concentration were fitted to the Michaelis-Menten kinetics to derive $K_M$ and $V_{max}$. $k_{cat}$ was derived from $V_{max}$ using the relationship $V_{max}=k_{cat}*[E_t]$. All graphs and curve fittings were processed using GraphPad Prism 9.

*Fluorometric coupled enzyme activity assay for NDST1 and mutants*

For the determination of pseudo-first order NDST1 kinetics with respect to polysaccharide substrate, varying concentrations of K5 or HS polysaccharide (low sulfated fraction; Iduron, GAG HS-I) were incubated with 250 nM NDST1, 350 nM SULT1A1-K65E-R68G, 20 µM PAPS and 4 mM 4MU-sulfate in 50 mM Tris-HCl pH 7.5, 15 mM $MgCl_2$, 1 mM DTT. Controls omitting polysaccharide were included and subtracted from the remaining reactions prior to further analysis.

For determination of pseudo-first order kinetics with respect to PAPS, a 2-fold serial dilution of PAPS from 50 µM was incubated with 0.5 $mg.mL^{-1}$ HS, 250 nM NDST1, 350 nM SULT1A1-K65E-R68G and 4 mM 4MU-sulfate in 50 mM Tris-HCl pH 7.5, 15 mM $MgCl_2$, 1 mM DTT. Controls omitting PAPS were included and subtracted from the remaining reactions prior to further analysis.

Evaluation of NDST1 mutants was performed using 1 µM enzyme in the presence of 350 nM SULT1A1-K65E-R68G, 20 µM PAPS, 0.5 mg.mL$^{-1}$ K5 and 4 mM 4MU-sulfate in 50 mM Tris-HCl pH 7.5, 15 mM $MgCl_2$, 1 mM DTT.

For evaluation of NDST1 activity in the presence of nanobodies, reactions were performed with an 8 point, 3-fold dilution of each nanobody from 90 µM in the presence of 250 nM NDST1, 350 nM SULT1A1-K65E-R68G, 20 µM PAPS, 0.5 mg.mL$^{-1}$ K5 and 4 mM 4MU-sulfate in 50 mM Tris-HCl pH 7.5, 15 mM $MgCl_2$, 1 mM DTT. NDST1 inhibition by ethylenediaminetetraacetic acid (EDTA) or diphenylamine (DPA) was demonstrated at 20 mM of either ligand under the same reaction conditions. Controls omitting polysaccharide were included and subtracted from the remaining reactions prior to further analysis.

The effect of NDST1 kinetics parameters in the presence of nAb7 or nAb13 was evaluated by addition of 30 µM nanobody to pseudo-first order NDST1 kinetics reactions with respect to K5 (see above). Reactions were incubated with 250 nM NDST1, 350 nM SULT1A1-K65E-R68G, 20 µM PAPS and 4 mM 4MU-sulfate in 50 mM Tris-HCl pH 7.5, 15 mM $MgCl_2$, 1 mM DTT. Controls omitting polysaccharide were included and subtracted from the remaining reactions prior to further analysis.

The effect of different metal ions on NDST1 activity was evaluated in the presence of 1 µM NDST1, 350 nM SULT1A1-K65E-R68G, 20 µM PAPS, 0.5 mg.mL$^{-1}$ K5 and 4 mM 4MU-sulfate in 50 mM Tris, pH 7.4, where the reaction buffer had been supplemented with 50 mM of either $MgCl_2$, $MnCl_2$, $ZnCl_2$, $CuCl_2$, $CaCl_2$ or $H_2O$ (metal free control). Control reactions containing 50 mM of either $MgCl_2$, $MnCl_2$, $ZnCl_2$, $CuCl_2$, $CaCl_2$ or $H_2O$ (metal free control), in the presence of 350 nM SULT1A1-K65E-R68G, 20 µM PAP and 4 mM 4MU-sulfate in 50 mM Tris, pH 7.4, were also included to assess the effect on SULT1A1-K65E-R68G activity.

In all cases reactions were performed in black half-area 96 well plates (Greiner) with a total reaction volume of 20 µL. All reactions were initiated by the addition of polysaccharide before recording the change in fluorescence with a Clariostar plate reader (BMG Labtech) using the 4MU preset. An 8 point 2-fold serial dilution standard curve of 4MU (from 10 µM) in 50 mM Tris-HCl pH 7, 15 mM $MgCl_2$, was used to calculate the evolution of 4MU during the reaction timecourse. For most datasets, initial reaction velocities at each substrate concentration were fitted to the Michaelis-Menten kinetics to derive $K_M$ and $V_{max}$ values. $k_{cat}$ was derived from $V_{max}$ using the relationship $V_{max}=k_{cat}*[E_t]$. For kinetics with the NDST1-nAb7 complex (**Figure 4f**) where initial reaction velocities did not plateau (i.e. $[S] << K_M$), a combined $k_{cat}/K_M$ value was derived from the slope of a linear fit to the data. $K_i$ values were calculated using the Morrison quadratic equation, which applies when $K_I$ and $[E_t]$ are in similar ranges:

$Y=Vo*(1-((((E_t+X+Q)-(((E_t+X+Q)^2)-4*E_t*X)^0.5))/(2*E_t)))$; $Q=(Ki*(1+([S]/K_M)))$. Graphs and curve fittings were processed using GraphPad Prism 9.

*Δ-Disaccharide analysis of time course-NDST1 treated K5*

600 µg of K5 was incubated with, 250 µM PAPS and 250 nM NDST1 in 50 mM Tris-HCl pH 7.4, 15 mM $MgCl_2$ for 18 hours at 37°C with shaking. Aliquots of the digest reaction were taken at timepoints, and heat inactivated at 95°C for 10 minutes, before being stored at -20°C until analysis.

Heparin lyase II (2 ug) was added to each sample and incubated at 30°C for a total of 24 hours, with the addition a further 2 ug of Heparin lyase II after 8 hours. Samples were heat denatured at 95°C for 5 minutes and stored at -20°C before analysis. Chromatographic separation of Heparin lyase II digested samples was performed using high performance anion exchange chromatography (HPAEC). Samples were made up to 1 mL in HPLC-grade $H_2O$ (Fisher) prior to being injected onto a ProPac PA-1 analytical column (4x250 mm), pre-equilibrated in HPLC-grade $H_2O$, at a flow rate of 1 $mL.min^{-1}$. The column was held under isocratic flow for 10 minutes, followed by elution of Δ-disaccharides using a linear gradient of NaCl (from 0 to 2 M NaCl in HPLC-grade $H_2O$) over 60 minutes. Elution was monitored by in-line UV detection of $A_{232}$ via the C=C unsaturated bond, introduced by Heparin lyase digestion. Retention times were compared to Δ-disaccharide reference standards (Iduron). The column was washed extensively with 2M NaCl and HPLC-grade $H_2O$ in between runs.

*Biotinylation of K5 polysaccharide*

K5 polysaccharide (4mM) was biotinylated at the reducing end as previously described[4,5] through reaction with 1:1 molar ratio of N-(aminooxyacetyl)-N'-(D-Biotinoyl) hydrazine in the presence of 100 mM aniline, in 100 mM acetate buffer pH 4.6. The reaction was incubated for 48 hours at 37°C before free biotin was removed by desalting on a C 10/40 column packed with Sephadex G25 (Cytiva), which had been pre-equilibrated in $dH_2O$. Separation was conducted over 1.2 CV with inline UV monitoring at 232 and 210 nm. Fractions eluting in the void volume corresponding to K5 were pooled and lyophilised. Biotinylation was assessed by dot blot; 2 µL of 1 $mg.mL^{-1}$ K5 or biotinylated K5 was spotted onto a nitrocellulose membrane and allowed to dry, before being blocked with 5% (w/v) BSA in PBS-T (PBS, 0.05% Tween-20) for 1 hour. The blocked nitrocellulose membrane was washed 3x with PBS-T before being incubated with 1:2000 HRP-streptavidin in 1% (w/v) BSA in PBS-T for 30 minutes. Following three washes with PBS-T, the presence of biotin was detected by the addition of SuperSignal West Femto chemiluminescence substrate (Thermo), and imaged using a ChemiDoc system (BioRad)

*Immunisation and library generation*

Antibodies to NDST1 were raised in a llama by intra-muscular immunization with purified protein using Gerbu LQ#3000 as the adjuvant. Immunisations and handling of the llama were performed under the authority of the project license PA1FB163A (University of Reading, UK). Total RNA was extracted from peripheral blood mononuclear cells, and $V_{HH}$ complementary DNAs were generated by RT-PCR. The pool of $V_{HH}$-encoding sequences was amplified by two rounds of nested PCR and cloned into the SfiI sites of the phagemid vector pADL-23c as previously described[6]. Electrocompetent E. coli TG1 cells were trans-formed with the recombinant pADL-23c vector, and the resulting TG1 phagemid library stock stored at -80 °C.

*Nanobody panning*

The VHH-displaying phage library was recovered by inoculation into 2x TY media supplemented with 100 µg.mL$^{-1}$ ampicillin, followed by incubation at 37°C, 250 rpm until the $OD_{600}$ reached 0.4–0.6. The culture was then infected with M13 helper phage and incubated for an hour at 37°C, before centrifugation at 2,800 g for 10 minutes at ambient temperature. The pellet was resuspended in 2x TY media supplemented with 100 µg.mL$^{-1}$ ampicillin and 50 µg.mL$^{-1}$ kanamycin and grown overnight at 25°C, 250 rpm with shaking to amplify the library. The amplified VHH-phage library was recovered by centrifugation at 3,200 g for 15 minutes at 4°C, to remove bacterial cells, and the supernatant collected. VHH-presenting phage were subsequently precipitated from the supernatant by the addition of 0.2 volumes of 20% (w/v) PEG6000, 2.5M NaCl and incubated on ice for one hour. The precipitated VHH-phage library was collected by centrifugation at 2,300 g for 10 minutes at 4°C, before being resuspended in 1 mL of ice-cold PBS. A further centrifugation step was performed at 20,000 g, 4°C for one minute, to remove any bacterial contaminants. The VHH-phage library was then reprecipitated by the addition of 0.2 volumes of 20% PEG6000 (w/v), 2.5 M NaCl and incubated on ice for 30 minutes, harvested by centrifugation at 20,000 g, 4°C for 15 minutes and resuspended in ice cold PBS, before being stored for a maximum of four weeks at 4°C.

The VHH-phage library was enriched for NDST1 binding VHHs by two rounds of bio-panning. The phage library was first blocked with StartingBlock (Thermo) for 30 minutes, followed by an incubation with 50 nM biotinylated NDST1 for an hour, rotating at room temperature. Pre-blocked Dyna beads (StartingBlock; Thermo Fisher Scientific) were mixed with the VHH-phage-biotinylated NDST1 mixture, and incubated for a further 15 minutes at room temperature. Dyna beads were captured and washed six times with PBS-T (1X PBS, 0.05% Tween-20), followed by one wash with PBS, before bound phage were released by digestion with 0.25 mg.mL$^{-1}$ trypsin (Sigma-Aldrich), in 10 mM Tris, 137 mM NaCl, 1 mM $CaCl_2$, for 30 minutes at room temperature. The eluted enriched VHH-phage library was collected and amplified by inoculation into exponentially growing TG1 *E.coli* cells, which were grown for 30 minutes at 37°C with shaking, then harvested by centrifugation at 2,800 g for 10 minutes before being resuspended in 1 mL of 2xTY media and plated onto LB agar containing 100 µg.mL$^{-1}$ ampicillin. Following incubation overnight at 37°C, TG1 *E.coli* cells were collected in 2xTY media containing 25% glycerol and stored at -80°C. The amplified enriched phage library was recovered as described previously, before a further round of bio-panning was performed with 5 nM biotinylated NDST1. Enrichment after bio-panning was determined by plating a 10-fold serial dilution of the recovered cell culture on LB agar plates containing 100 µg.mL$^{-1}$ ampicillin.

For screening for NDST1 binding clones, 93 individual clones were selected and grown in 2x TY media (100 µg.mL$^{-1}$ ampicillin) overnight at 37°C, 250 rpm. Overnight cultures were inoculated into 2x TY media containing 100 µg.mL$^{-1}$ ampicillin and incubated for three hours at 37°C, 250 rpm shaking, before being infected with M13 helper phage. The infected cultures were grown for a further hour at 37°C, 250 rpm before being centrifuged at 2,800 g for 10 minutes at 4°C. The cell pellet was resuspended in 2x TY media containing 100 µg.mL$^{-1}$ ampicillin and 50 µg.mL$^{-1}$ kanamycin, before being grown at 25°C, 250 rpm overnight. Clonal VHH-presenting phage were harvested from the culture supernatant by centrifugation at 3,500 rpm for 15 min at 4°C , to remove bacterial cells, and stored at 4°C for further analysis.

*Enzyme-linked immunosorbent assays to identify NDST1 binding phage*

A 96-well high binding microtiter plate (Greiner) was coated with 5 µg.mL$^{-1}$ neutravidin in 100 uL PBS per well, and stored overnight at 4°C. The plate was then washed 5x with PBS-T before the addition of 100 µL of biotinylated NDST1 (50 nM in 20 mM MES, 100 mM NaCl, pH 6.5, 0.1% fat free skimmed milk) per well, and then incubated for one hour at room temperature with agitation. The plate was washed 5x with 20 mM MES, 100 mM NaCl, pH 6.5 before being blocked with PBS-T, 2% non-fat skimmed milk (250 µL per well) for one hour at room temperature, with agitation. Following a further 5 washes with PBS-T, 100 µL of each clonal VHH-presenting phage stock (diluted with 1 volume of PBS-T, 2% non-fat skimmed milk) was added to each well and incubated at room temperature, with agitation, for one hour. The plate was washed five times with PBS-T, before being incubated for one hour at room temperate with 100 µL αM13-HRP (1:5000 dilution in StartingBlock; Thermo) per well, with agitation. The plate was washed 5x with PBS-T before the addition of 100 µL per well TMB substrate (SeraCare). Absorbance was read at 405 nM using a ClarioStar plate reader.

*Nanobody expression and purification*

Identified VHH hits were cloned into the pOPINO vector and transformed into WK6 *E. coli* cells using electroporation. Pre-cultures of nanobodies were grown in 2xTY media supplemented with 100 µg.mL$^{-1}$ ampicillin and 2% (w/v) glucose at 37°C, 250 rpm shaking, overnight. 1L of TB media containing 100 µg.mL$^{-1}$ ampicillin, 0.1% glucose (w/v) and 1 mM $MgCl_2$ was inoculated with the preculture and grown at 37°C, 250 rpm until the OD600 reached 1–1.2. Expression was induced with 1 mM IPTG and the culture incubated at 28°C overnight, 250 rpm shaking. Cells containing the nanobody were collected by centrifugation at 5000 rpm for 15 minutes at 4°C. Nanobodies were extracted from the periplasm by resuspending the cell pellet in TES buffer (200 mM Tris pH 8.0, 0.5 mM EDTA, 500 mM sucrose) overnight with constant agitation. Two volumes of 50 mM Tris pH 8.0, 125 mM sucrose, supplemented with 120 kU DNase I was then added, and the resulting suspension incubated for a further 2 hours to lyse cells. Cell debris were removed by centrifugation at 16,000 rpm for 30 minutes at 4°C and the supernatant collected. The supernatant was diluted with 5 volumes of PBS, pH 8.0 and filtered before being loaded onto a 5 mL HisTrap FF column (Cytiva) pre-equilibrated in Histrap buffer A (PBS, 30 mM imidazole). The column was washed with 5 CV of the same buffer, before elution of the nanobody with a linear gradient of 0-100% Histrap buffer B (PBS, 300 mM imidazole) over 10 CV. Fractions containing the nanobody were pooled and concentrated using a 10 kDa MWCO Vivaspin filter. Nanobodies were purified further by SEC using a HiLoad 16/600 Superdex 75 pg column pre-equilibrated in PBS. Nanobody containing fractions were pooled, concentrated to >2 mg.mL$^{-1}$ and flash frozen in $LN_2$. Purified nanobodies were stored at -80°C until required.

*Binding analyses – SPR*

Surface plasmon resonance experiments were performed using a Biocore T200 system (GE Healthcare) primed with 20 mM MES pH 6.5, 100 mM NaCl, 0.05% Tween-20. All assays were performed using a BiotinCAP Sensor chip (GE Healthcare) at 25°C. To determine the binding affinity of nanobody clones for NDST1, biotinylated NDST1 (10 µM) was immobilised onto the sample channel of the sensor chip at a flow rate of 2 µL.min$^{-1}$ for 600 s. The reference channel was left blank. A titration of each nanobody (from 10 µM) was injected over both sensor channels at a flow rate of 30 µL.min$^{-1}$, for 100 s. Dissociation was monitored for 100 s. Steady state binding responses were fitted to a 1:1 binding model (One-site specific binding) using GraphPad prism 9 to calculate $K_D$ values. In combinatorial assays nanobodies were either pre-mixed (1:1, 10 µM) or injected alone at a flow rate of 30 µL.min$^{-1}$ for 100s, with monitoring of the dissociation for 100 s, followed by comparison of the maximal response during the association phase.

*Binding analyses – BLI*

Biolayer interferometry (BLI) experiments were performed with an Octet R8 system (Sartorius) using Octet SA biosensors. For determination of the binding affinity of nanobodies to NDST1, biosensors were hydrated with two washes of 20 mM MES pH 6.5, 100 mM NaCl, before immobilization of biotinylated NDST1 (75 nM) for 300 s in the same buffer. The loaded sensors were washed twice for 60 s in PBS-T, and the baseline recorded. Association of nanobodies in PBS-T to immobilized NDST1 or unloaded reference sensors were recorded for 100–600s, followed by measurement of the dissociation in PBS-T for 100–300s. All steps were performed with shaking at 1,000 rpm at 25°C. Control reactions containing no nanobody and reference sensors were subtracted prior to alignment and smoothing of the sensorgrams with Savitzky-Golay filtering using Octet Analysis studio (Sartorius). Steady state binding responses were fitted to a 1:1 binding model (One-site specific binding) using GraphPad prism 9 to calculate $K_D$ values.

For determination of the binding affinity of NDST1 to K5, Octet SA biosensors were hydrated with three washes in PBS-T for 300 s before immobilization of biotinylated-K5 (25 $\mu g.mL^{-1}$ or 50 $\mu g.mL^{-1}$) in the same buffer for 300 s. The loaded sensors were washed twice, for 60 s, in PBS-T and the baseline recorded. Association of NDST1 to immobilized biotinylated-K5 or unloaded reference sensors was recorded for 300 s followed by measurement of dissociation in PBS-T for 300s. All steps were performed with shaking at 1,000 rpm at 25°C. Control reactions containing no NDST1 and reference sensors were subtracted prior to smoothing of the sensorgrams with Savitzky-Golay filtering using Octet Analysis studio. Data were fitted using 2:1 heterogenous binding model with global fitting in Octet Analysis studio to determine $k_{assoc}$, $k_{dis}$ and $K_D$ values. The effect of nanobodies nAb7 and nAb13 on NDST1 binding to K5 was assessed using the same conditions, with the inclusion of 10 µM nanobody. To evaluate the effect of NaCl on NDST1 binding to K5, an 8 point 1.5-fold dilution of 1.5 M NaCl into 10 mM phosphate buffer pH 7.4, 0.05% Tween-20 was performed using the same assay conditions.

*Generating NDST1 nanobody complexes*

NDST1 and nanobodies were incubated in 20 mM MES pH 6.5, 200 mM NaCl, at 1:1 mass ratios, for 1 h at room temperature. Complexes were isolated by size exclusion chromatography on a Superdex 200 Increase 10/300 GL column (Cytiva), pre-equilibrated in the same buffer. NDST1-nAb7 and NDST1-nAb13 containing fractions were pooled and concentrated to >5 $mg.mL^{-1}$, using a 10 kDa MWCO vivaspin concentrator. Small aliquots were flash frozen using liquid nitrogen and stored at -80°C for further use, or used immediately.

*Cryo-EM sample preparation*

Frozen aliquots of NDST1-nAb7 or NDST1-nAb13 complex were thawed rapidly and protein concentration adjusted using 50 mM Tris-HCl pH 7.5, supplemented with PAP (5 mM) and $MgCl_2$ (15 mM). 2.5 µL of these samples were applied to Quantifoil R1.2/1.3 300 Cu mesh grids, glow discharged for 60 s at 30 mA negative current in a GloQube Plus glow discharger (Quorum Technologies). Grids were blotted for 1–3 s using a blot force of 3 before plunging into liquid ethane using a Vitrobot Mark IV (Thermo Fisher) set to 4°C and 100% humidity. Grids were clipped into Autogrid rings (Thermo Fisher) before loading into microscopes for screening and data collection.

*Cryo-EM data collection and processing*

Th NDST1 alone and NDST1-nAb13 datasets were collected from the same cryo-EM grid, in which a large proportion of NDST1-nAb13 complexes had dissociated, enabling classification of both nAb13 bound and unbound particles. Data was collected on a Krios G4 microscope at the Rosalind Franklin Institute, using an accelerating voltage of 300 kV, Falcon 4i direct electron detector and Selectris X energy filter with a slit width of 10 eV. Movies were collected using EPU in EER format at 165kx magnification, using a total dose of 50 electrons per $Å^2$, a calibrated pixel size of 0.73 Å, and target defocus values of -1.2 µm to -2.6 µm in 0.2 µm intervals.

The NDST1-nAb7 dataset was collected on a Krios G3i microscope at the Oxford Particle Imaging Centre (OPIC), using an accelerating voltage of 300 kV, Falcon 4 direct electron detector and Selectris X energy filter with a silt width of 10 eV. Movies were collected using EPU in EER format at 165kx magnification, using a total dose of 50 electrons per $Å^2$, a calibrated pixel size of 0.7303 Å, and target defocus values of -1.2 µm to -2.6 µm in 0.2 µm intervals.

EM processing workflows for each dataset are outlined in **Figure S6**. In brief, micrograph pre-processing was performed in cryoSPARC v2.14.1[7-9] live interface using path-motion CTF correction. Initial particles were picked using unbiased blob-picking, and good particles isolated via two rounds of 2D classification. 2D class averages with secondary structure features were used to guide template-based particle picking, followed by further rounds of 2D classification. *Ab initio* models generated in cryoSPARC were used as non-biased references for 1 or more rounds of heterogenous refinement.

The best classes from heterogenous refinement, as judged by resolution, were pooled and refined further using cryoSPARC non-uniform refinement, followed by rounds of local and global CTF refinement where appropriate, followed again by non-uniform refinement. Final postprocessing and local resolution estimates were carried out in Relion 3.1.1[10].

3D-variability analyses on the NDST1-nAb7 and NDST1-nAb13 complexes were each carried out using the cryoSPARC pipeline, incorporating 3 variability components. Focussed local refinements for both complexes were carried out in cryoSPARC, with masking to subtract either the NTD or the sulfotransferase domain, generating subtracted particles for the deacetylase-sulfotransferase-nAb or NTD-deacetylase-nAb regions respectively. Locally refined volumes were postprocessed using Relion, before being combined for model building purposes using UCSF ChimeraX, by taking the maximum value from either locally refined map at each voxel. All other analyses and depositions were carried out with the individual locally refined maps.

For all structures, cryo-EM model building was carried out in COOT[11], using starting models initially derived from AlphaFold2 (NDST1) or the PDB (for nAb7, from accession 7TGF). Models were improved by manual building using COOT, iterated with rounds of real-space refinement using PHENIX.REFINE[12]. The nAb13 nanobody structure was derived from the nAb7 structure by conversion to an all Cα chain in COOT, and removal of the long nAb7 unique CDR3 loop (R98–F116), before fitting the Cα model into the NDST1-nAb13 volume by 5 rounds of rigid-body refinement in PHENIX.REFINE. Structural figures were generated using ChimeraX[13]. Electrostatic surfaces were calculated using ChimeraX. Binding interactions were determined using PISA analysis[14] with default settings. RMSD values were calculated using CCP4MG[15]. Sequence alignment values were calculated using ClustalΩ[1]. All cryo-EM processing and model building statistics are shown in **Table S1**.

*Molecular Docking*

Interactions between NDST1 and HS were analysed by molecular docking using GlycoTorch Tools (GTV) and GlycoTorch Vina[16]. The protonation of residues of both protein and ligands were set to reflect physiological pH. Glycam carbohydrate builder (Glycam.org) was used to build the two octasaccharides, $[GlcNAc\text{-}GlcA]_4$ and $[GlcNAc\text{-}GlcA]_2\text{-}[GlcNS\text{-}GlcA]_2$. GTV was used to convert the octasaccharides obtained from Glycam to pdbqt format. A centroid selected from the PAP binding site and metal binding sites from the cryo-EM structure was used to define the coordinates for the docking of heparosan oligosaccharides. The docking the oligosaccharides was performed using the grid box sizes of 40 × 50 × 40 Å and 55 × 40 × 25 Å from the centroids in the deacetylase and sulfotransferase domains respectively. The octasaccharides were docked using GlycoTorch Vina with energy range, exhaustiveness and num_modes values set to be 12, 80, and 100, respectively. The conformations chosen for further analysis were based on the best-estimated affinities and their ranking in the clustering process. Docking visualization and figures were generated using ChimeraX[13].

*Inductively coupled plasma optical emission spectrometry (ICP-OES)*

Transferrin or apotransferrin (440 mg.mL$^{-1}$) was digested with the addition of 0.25 volumes of $HNO_3$ and diluted two-fold across 5 concentrations in 1.25% HNO3. 150 mg.mL$^{-1}$ NDST1 was by the addition of 0.25 volumes of $HNO_3$ in a total volume of 4 mL. Following digestion, each sample was filtered using a 0.25 mm PVDF filter prior to measurement of Ca ($\lambda_{em}$ = 422.673 nm), Co ($\lambda_{em}$ = 238.892 nm), Cu ($\lambda_{em}$ = 324.754 nm), Fe ($\lambda_{em}$ = 260.709 nm), Mg ($\lambda_{em}$ = 279.553 nm), Mn ($\lambda_{em}$ = 279.482 nm) and Zn ($\lambda_{em}$ = 206.2 nm) using an Agilent Technologies 700 Series ICP-OES. ARISTAR® multi-element (28) quality control standard (VWR), containing the aforementioned metal ions at a concentration of 1 mg.mL$^{-1}$ each, was serially diluted 2-fold across 5 concentrations covering the range of 0.0625 - 1 ppm to allow the construction of a calibration curve for each element. Suitable wavelengths were chosen where interferences were not observed, and a sufficient signal:noise ratio was observed above baseline. All samples were obtained as a mean of 3 replicates under a plasma argon flow rate of 15 L.min$^{-1}$, auxiliary argon flow rate of 1.5 L.min$^{-1}$ and a nebulizer argon flow rate of 0.75 L.min$^{-1}$.

**Methods references**


1. Sievers, F. et al. Fast, scalable generation of high-quality protein multiple sequence alignments using Clustal Omega. *Mol Syst Biol* **7**, 539 (2011).
2. Liu, H. & Naismith, J.H. An efficient one-step site-directed deletion, insertion, single and multiple-site plasmid mutagenesis protocol. *BMC Biotechnol* **8**, 91 (2008).
3. Lu, L.Y., Hsu, Y.C. & Yang, Y.S. Spectrofluorometric assay for monoamine-preferring phenol sulfotransferase (SULT1A3). *Anal Biochem* **404**, 241-3 (2010).
4. Thakar, D. et al. A quartz crystal microbalance method to study the terminal functionalization of glycosaminoglycans. *Chem Commun (Camb)* **50**, 15148-51 (2014).
5. Su, D. et al. Analysis of protein-heparin interactions using a portable SPR instrument. *PeerJ Analytical Chemistry* **4**, e15 (2022).
6. Huo, J. et al. A potent SARS-CoV-2 neutralising nanobody shows therapeutic efficacy in the Syrian golden hamster model of COVID-19. *Nat Commun* **12**, 5469 (2021).
7. Punjani, A., Zhang, H. & Fleet, D.J. Non-uniform refinement: adaptive regularization improves single-particle cryo-EM reconstruction. *Nat Methods* **17**, 1214-1221 (2020).
8. Rubinstein, J.L. & Brubaker, M.A. Alignment of cryo-EM movies of individual particles by optimization of image translations. *J Struct Biol* **192**, 188-95 (2015).
9. Punjani, A., Rubinstein, J.L., Fleet, D.J. & Brubaker, M.A. cryoSPARC: algorithms for rapid unsupervised cryo-EM structure determination. *Nat Methods* **14**, 290-296 (2017).
10. Zivanov, J. et al. New tools for automated high-resolution cryo-EM structure determination in RELION-3. *Elife* **7**(2018).
11. Casanal, A., Lohkamp, B. & Emsley, P. Current developments in Coot for macromolecular model building of Electron Cryo-microscopy and Crystallographic Data. *Protein Sci* **29**, 1069-1078 (2020).
12. Liebschner, D. et al. Macromolecular structure determination using X-rays, neutrons and electrons: recent developments in Phenix. *Acta Crystallogr D Struct Biol* **75**, 861-877 (2019).
13. Pettersen, E.F. et al. UCSF ChimeraX: Structure visualization for researchers, educators, and developers. *Protein Sci* **30**, 70-82 (2021).
14. Krissinel, E. & Henrick, K. Inference of macromolecular assemblies from crystalline state. *J Mol Biol* **372**, 774-97 (2007).
15. McNicholas, S., Potterton, E., Wilson, K.S. & Noble, M.E. Presenting your structures: the CCP4mg molecular-graphics software. *Acta Crystallogr D Biol Crystallogr* **67**, 386-94 (2011).
16. Boittier, E.D., Burns, J.M., Gandhi, N.S. & Ferro, V. GlycoTorch Vina: Docking Designed and Tested for Glycosaminoglycans. *J Chem Inf Model* **60**, 6328-6343 (2020).
